# Multiplexed imaging of immune cells in staged multiple sclerosis lesions by mass cytometry

**DOI:** 10.1101/638015

**Authors:** Valeria Ramaglia, Salma Sheikh-Mohamed, Karen Legg, Olga L Rojas, Stephanie Zandee, Fred Fu, Olga Ornatsky, Eric C Swanson, Alexandre Prat, Trevor D McKee, Jennifer L Gommerman

## Abstract

Multiple Sclerosis (MS) is characterized by demyelinated and inflammatory lesions in the brain and spinal cord. Lesions contain immune cells with variable phenotypes and functions. Here we use imaging mass cytometry (IMC) to enable the simultaneous imaging of 15+ proteins within 11 staged MS lesions. Using this approach, we demonstrated that the majority of demyelinating macrophage-like cells in active lesions were derived from the resident microglial pool. Although CD8^+^ T cells predominantly infiltrated the lesions, CD4^+^ T cells were also abundant but localized closer to blood vessels. B cells with a predominant switched memory phenotype were enriched across all lesion stages and were found to preferentially infiltrate the tissue as compared to unswitched B cells which localized to the vasculature. We propose that IMC will enable a comprehensive analysis of single-cell phenotypes, their functional states and cell-cell interactions in relation to lesion morphometry and demyelinating activity in the MS brain.

## INTRODUCTION

Multiple sclerosis (MS) is a disease with profound heterogeneity in the neuropathological and immunopathological appearance of lesions in the central nervous system (CNS)[28]. Recent consensus has standardized staging of MS brain tissue into categories including normal-appearing white matter (NAWM), (p)reactive lesions (or “pre-phagocytic” lesions)[3] which may represent an initial lesion[32,1], periplaque white matter (PPWM) which is immediately adjacent to a lesion, early or late active demyelinating lesions, mixed active/inactive demyelinating lesions (also called slowly expanding or “smouldering”[16]), and inactive lesions[25]. The pattern of demyelination can also be fundamentally different between patients, with pattern I being T cell-mediated, pattern II being IgG- and complement-mediated, and pattern III and IV characterized by a primary oligodendrocyte dystrophy reminiscent of virus- or toxin-induced demyelination rather than autoimmunity[15].

Lymphocytes, microglia and macrophages are associated with active demyelination and neurodegeneration in the MS brain[18,15] and are thought to play key roles in the disease process, as supported by studies in experimental models (reviewed in[36]). Depending on the type of lesion and the sub-region within a lesion (for example center vs edge), different myeloid and lymphoid cells can be found. These have a variety of phenotypes that reflect activation state and pathologic potential. With respect to myeloid cells, yolk sac-derived (resident) microglia and blood-derived (recruited) monocytes/macrophages accumulate at sites of active demyelination and neuroaxonal injury[15]. Microglia and macrophages within the MS brain can lose their normally homeostatic properties and acquire a pro-inflammatory phenotype with expression of molecules involved in phagocytosis, oxidative injury, antigen presentation and T cell co-stimulation[17]. Either via T cell-mediated recognition of myelin epitopes[41] or complement binding to myelin autoantibodies[40], these lymphocyte-dependent events initiate a process that results in the activation of microglia and recruitment of macrophages at the lesion site. Microglia and macrophages become activated and internalize myelin, degrading it within their lysosomes. The detection of small (myelin oligodendrocyte glycoprotein, MOG) or large (proteolipid protein, PLP) myelin proteins indicates the temporal development of myelin destruction[28,25].

In terms of lymphocytes, MS lesions contain T cells and CD20^+^ B cells[31]. In active lesions, CD8^+^ T cells proliferate and have an activated cytotoxic phenotype. Subsequently, some CD8^+^ T cells are destroyed by apoptosis while others, with tissue-resident memory features, persist. Tissue resident memory T cells lose expression of surface molecules that are involved in the egress of leukocytes from inflamed tissue, which has been suggested as a potential mechanism responsible for the compartmentalized inflammatory response in established lesions[31]. CD4^+^ T cells are also found in MS lesions and have been shown to produce cytokines such as IL-17 and IFN*γ*[23]. B cells are thought to differentiate into plasma cells, perhaps *in situ*, but little is known about their phenotype[31].

While past and recent immunohistological studies have provided insigths into the types of immune cells populating MS lesions at different lesional stages and the neurodegenerative changes that accompany these infiltrating immune cells[31,29,44,35,14,15], this type of analysis requires immunohistological staining of serial sections and is limited to the number of analytes that can be simultaneously visualized on a given tissue section. Thus, a comprehensive analysis of single-cell phenotypes and functional states in relation to demyelination within MS tissue is lacking. To circumvent this challenge, we have employed imaging mass cytometry (IMC). IMC uses time-of-flight inductively coupled plasma mass spectrometry to detect dozens of markers simultaneously on a single tissue section. It achieves this by measuring the abundance of metal isotopes tagged to antibodies and indexed against their source location[10]. Applying this new technology to post-mortem MS brain tissue, we carefully analysed staged lesions in a case with severe rebound MS disease activity after natalizumab (NTZ) cessation[26]. The data we collected suggests that imaging mass cytometry, in combination with existing imaging techniques, can profoundly impact our knowledge of the inflammatory response and tissue injury in the MS brain.

## MATERIALS & METHODS

### Patient case report and pathological analysis of the brain

The clinical and pathological characteristics of the case reported in this study have been previously published[26]. Briefly, the patient was a 32-year-old female, diagnosed with relapsing-remitting MS in 2005. Natalizumab (NTZ) therapy was intitiated (expanded disability status scale (EDSS) 5.0 per relapse, and 3.5 upon induction on NTZ) but stopped after 2 years because, although clinically and radiologically stable (EDSS 2.0), the patient tested positive to the John Cunningham virus (JVC) virus antibody titer. Glatiramer acetate was started 1 month prior to NTZ cessation, and the patient received a 5-day course of intravenous (iv) methylprednisolone after the last NTZ infusion. Four months later, the patient was hospitalized for the presentation of new motor and cognitive deficits. Over the course of 2 weeks, the patient worsened (EDSS 9.5). Despite daily course of iv methylprednisolone, the patient developed several new gadolinium-enhancing lesions on repeated MRI. Since no clinical or radiological improvements were observed, the family decided to stop active care, as per patient’s previous wishes. Autopsy was performed within 1 hour post-mortem, 4 days after withdrawal of all medication.

The patient had previously provided written consent for post-mortem donation of the CNS to research (ethics committee approval number BH.07.001). Pathological analysis of the brain revealed abundant, active demyelinating, and highly inflammatory MS lesions with immunological pattern II (IgG- and complement-mediated)[26], according to Lucchinetti et al[28]. Despite the extent of the inflammation, progressive multifocal leukoencephalopathy or immune reconstitution inflammatory syndrome (IRIS) were excluded, from a pathological point of view, although a later study proposed that Epstein-Barr virus-associated IRIS could have been the possible cause of the fulminant MS relapse in this case[39]. Of note, immunohistochemistry and qPCR for JCV was reported negative. Finally, the neuropathologist-confirmed diagnosis of severe MS rebound inflammatory demyelinating activity after NTZ withdrawal[26]. The non-neurological control case was an 86-year-old female that died of cardiac arrest. This control case was obtained from the Netherlands Brain Bank (VU Medical Center ethic committee approval Reference number 2009/148).

### Sample characterization

Our study was performed on two frozen tissue blocks from the MS case and one frozen tissue block from the non-neurological control case. We analysed immunopathological changes in the white matter of the MS case, focusing on the following regions of interest (ROI) in lesions staged acconding to Kuhlmann et al[25] (see **Figure 1 – Figure Supplement 1**): normal-appearing white matter (NAWM), located > 1 cm distant from any lesions (detected in the block); the periplaque white matter (PPWM), located < 1 cm distant from any lesions; (p)reactive lesions, that may represent the initial stage of a lesion[32,1,25], defined by the presence of microglia/macrophages in the absence of (obvious) demyelination, as described by Luchetti et al[29]; early active demyelinating lesions defined by presence of microglia/macrophages with early (MOG) and late (PLP) myelin degradation products throughout the lesion, as described[7], and previously shown for this MS case[26], supporting the abundance of demyelinating activity in this type of lesions; the active edge of mixed active-inactive demyelinating lesions defined as slowly expanding lesions or smouldering lesions by Frischer et al, that are normally present in the progressive stage of MS[16] but have also been described in cases with acute MS[44]; and lastly the inactive center of a mixed active-inactive demyelinating lesion. The normal white matter of control (WMC) was analysed as a reference background for the immunopathological composition of the lesions in the MS case. The lesion types and ROI analysed are indicated in **Table 1**.

**Table 1.** Lesion types and regions of interest.

| **Case** | **Tissue Block**<br>**(anatomical location)** | **Lesion Type** | **Region of interest** |
| --- | --- | --- | --- |
| C,<br>95-056 | A<br>(superior frontal gyrus) |  | WMC |
| MS,<br>AB129 | CL3a<br>(cerebellum) |  | NAWM |
|  |  | 2x Active demyelinating (pattern II) | PPWM, center |
|  |  | 3x Mixed active/inactive demyelinating | PPWM, edge, core |
|  | CR4a<br>(cerebellum) |  | NAWM |
|  |  | (p)reactive |  |
|  |  | 3x Active demyelinating (pattern II) | center |
|  |  | 2x Mixed active/inactive demyelinating | edge, core |

C, control; MS, multiple sclerosis; WMC, white matter of control; NAWM, normal-appearing white matter; PPWM, periplaque white matter.

### Selection of inflammatory markers

To define inflammatoty cells as a whole, we used CD45, a general marker for microglia, macrophages and lymphocytes, with CD45^low/+^ indicative of microglia and CD45^high^ indicative of macrophages and lymphocytes. For microglia we additionally used TMEM119, that is expressed on yolk sac-derived (resident) microglia but not on recruited blood-derived macrophages[5,38]. Other markers were used to detect phagocytosis (CD68) and capacity for antigen presentation (human leukocyte antigen, HLA). All T cells were detected with the cellular marker CD3. CD8α detected MHC class I restricted T cells while CD4 detected MHC class II restricted T cells. All B cells were identified by the expression of either the kappa (κ) or lambda (λ) allelic variants of the immunoglobulin light chain. CD38 was used to detect a multifunctional molecule expressed by leucocytes in general and involved in the activation of T cells and B cells. IgM was used to identify naïve and non-class switch memory B cells, in addition to detecting free immunoglobulins. Furthermore, microglia and T cells express the transcription factor nuclear factor of activated T cells (NFAT1), that translocates to the nucleus upon activation[12]. Therefore, we determined the localization of NFAT1 as an additional activation antigen of CD45^low/+^TMEM119^+^ microglia and CD45^high^CD3^+^CD8α^+^ or CD45^high^CD3^+^CD8α^−^ T cells. We also used the Ki67 marker of cell proliferation and PLP to identify myelin. Blood vessels were identified using markers of extracellular matrix (collagen) and endothelial cells (CD31). Each antibody clone was first titrated for immunofluorescence staining in control and MS tissue, according to the dilutions shown in **Table 2**, prior to methial-conjugation and IMC application.

**Table 2.** Antibodies used for Imaging Mass Cytometry.

| **Antibody** | **Target** | **Metal Tag** | **Antibody Clone** | **Concentration (µg/ml)** | **Source (Catalogue)** |
| --- | --- | --- | --- | --- | --- |
| **Ir-Intercalator** | DNA | Ir191<br>Ir193 |  | 0.5 | Fluidigm (201192A) |
| **PLP (proteolipid protein)** | Myelin | 141Pr | Plpc1 | 20, 40<sup>a</sup>, 10<sup>b</sup> | Bio-Rad (MCA839G) |
| **CD38** | Leukocytes | 167Er | HIT2 | 0.5 | Fluidigm (3167001B) |
| **CD45** | Leukocytes/<br>Macrophages | 154Sm | HI30 | 0.5 | Fluidigm (3154001B) |
| **CD68** | Lysosomal Marker (phagocytic cells) | 159Tb | KP1 | 10 | Fluidigm (3159035D) |
| **HLA (human leukocyte antigen)** | Microglia/<br>Macrophages | 147Sm | LN3 | 5, 10<sup>a</sup>, 5<sup>b</sup> | Abcam (Ab55152) |
| **TMEM119** | Microglia | 155Gd | Polyclonal | 10, 5<sup>a</sup> | Sigma-Aldrich (HPA051870) |
| **CD3** | T Cells | 170Er | UCHT1 | 10 | Fluidigm (3170001B) |
| **CD4** | CD4<sup>+</sup> T Cells | 176Yb | SK3 | 5, 25<sup>a</sup> | BioLegend (344602) |
| **CD8α** | CD8<sup>+</sup> T cells | 162Dy | RPA-T8 | 10 | Fluidigm (3162015B) |
| **Granzyme B** | CD8<sup>+</sup> T Cell Activation | 171Yb | GB11 | 20, 50<sup>a</sup> | ThermoFisher Scientific (MA1-80734) |
| **Ig Kappa** | Immunoglobulin (light chain) | 160Gd | MHK-49 | 0.33 | Fluidigm (3160006B) |
| **Ig Lambda** | Immunoglobulin (light chain) | 151Eu | MHL-38 | 0.33 | Fluidigm (3151004B) |
| **IgM** | Immunoglobulin | 172Yb | MHM-88 | 2 | Fluidigm (3172004B) |
| **Collagen** | Blood Vessels | 169Tm | Polyclonal | 0.25 | Fluidigm (3169023D) |
| **CD31** | Endothelial cells | 145Nd | C31.3+<br>C31.7+<br>C31.10 | 10 | LSBio (LS-C390863) |
| **NFAT1 (Nuclear Factor of activated T cells)** | Cell activation | 143Nd | D43B1 | 20 | Fluidigm (3143023A) |
| **Ki67** | Cell Proliferation | 168Er | Ki-67 | 10 | Fluidigm (3168001B) |

<sup>a</sup>, <sup>b</sup>concentration of the unlabeled antibody by immunofluorescence<sup>a</sup> or immunohistochemistry<sup>b</sup>

### Histology, immunohistochemistry (IHC) and immunofluorescence (IF)

Ten-micron frozen tissue sections were mounted on Superfrost Plus glass slides (Knittel Glass) and stored at −80°C until they were stained.

#### Histology

On the day of the staining, the slides were brought at room temperature and post-fixed in 10% formalin for 3 hours. Tissue sections were stained with Hematoxylin and Eosin (HE)/Luxol Fast Blue (LFB) to detect myelin lipids and Oil red O (ORO) to detect neutral lipids in phagocytic macrophages, as previously published[34].

#### IHC

On the day of the staining, the slides were brought to room temperature and post-fixed in ice-cold acetone for 10 minutes. Myelin protein was detected using an antibody for proteolipid protein (PLP) and microglia/macrophages were detected using an antibody for human leukocyte antigen (HLA). Endogenous peroxidases activity was blocked by incubation in PBS with 0.3% H_2_O_2_ for 20 minutes at room temperature. Non-specific protein binding was blocked by incubation with 10% normal goat serum (DAKO). Primary antibodies were applied overnight at 4°C, diluted in normal antibody diluent (Immunologic, Duiven, The Netherlands) according to dilutions noted in **Table 2**. The following day, sections were incubated with a post-antibody blocking solution for monoclonal antibodies (Immunologic) diluted 1:1 in PBS for 15 minutes at RT. Detection was performed by incubating tissue sections in secondary Poly-HRP (horseradish peroxidase)-goat anti-mouse/rabbit/rat IgG (Immunologic) antibodies diluted 1:1 in PBS for 30 minutes at RT followed by application of DAB (3,3-diaminobenzidine tetrahydrochloride (Vector Laboratories, Burlingame, CA, U.S.A.) as a chromogen. Counterstaining was performed with hematoxylin (Sigma-Aldrich) for 10 minutes. The sections were subsequently dehydrated through a series of ethyl alcohol solutions and then placed in xylene before being coverslipped with Entellan mounting media (Sigma Aldrich). The colorimetric staining was visualized under a light microscope (Axioscope, Zeiss), connected to a digital camera (AxioCam MRc, Zeiss) and imaged with Zen pro 2.0 imaging software (Zeiss).

#### IF

On the day of the staining, the slides were brought to room temperature and incubated in ice-cold acetone for 10 minutes followed by 70% ethanol for 10 minutes to reduce the autofluorescence signal derived from the fatty myelin sheets. Slides were subsequently rehydrated in 0.05% PBS-tween for 10 minutes at room temperature followed by incubation in 10% normal goat serum (DAKO) to block nonspecific binding sites. Sections were then incubated overnight at 4 with primary antibody diluted in 3% normal goat serum (see dilutions in **Table 2**). Primary antibodies were detected using fluorochrome-conjugated secondary antibodies (Sigma-Aldrich) diluted 1:200 in 1% Triton-X100. Sections were incubated with DAPI (Sigma Aldrich) diluted 1:3000 to visualize the nuclei. Slides were washed in PBS, air dried and mounted in aqueous mounting medium. Using the appropriate filters, the IF signal was visualized with an Axio Imager Z1, Zeiss microscope connected to a digital camera (AxioCam 506 mono, Zeiss) and imaged with Zen pro 2.0 imaging software (Zeiss).

To control for antibodies specificity, tissue sections were stained according to the IF or IHC protocols described above except for the primary antibody incubation step, which was omitted.

### Imaging mass cytometry

The work flow of imaging mass cytometry is shown in **Figure 1 – Supplement Figure 2** and explained in detail below.

#### Conjugation of antibodies with lanthanide metals

Lanthanide metal-conjugated antibodies were either obtained from Fluidigm (Markham, Ontario, Canada) or conjugated at SickKids-UHN Flow and Mass Cytometry Facility (Toronto, Ontario, Canada), using the MaxPar X8 labelling kit from Fluidigm (catalogue number 201169B) as previously described[19]. Briefly, a purified carrier-free antibody was partially reduced with TCEP buffer (Fluidigm, catalogue number 77720) at 37°C. The reduced antibody was then incubated with an excess of metal-loaded MaxPar X8 polymer for 90 minutes at 37°C. The metal-labeled antibody was then recovered using a 50kDa size exclusion filter. The percent yield of metal-conjugated antibody was determined by measuring the absorbance of the conjugate at 280nm. The recovery of our metal-conjugated antibodies was 69-78%. Antibody stabilizer was then added to the metal-conjugated antibodies before long-term storage at 4°C.

#### Staining for Imaging Mass Cytometry

On the day of staining, the slides were brought to room temperature and rehydrated with 0.05% PBS-Tween in a humidified chamber for 20 minutes at room temperature. Non-specific protein binding was blocked by incubation with 10% normal goat serum for 20 minutes at room temperature followed by incubation with blocking solution (ThermoScientific Superblock Blocking Buffer in PBS) for 45 minutes at room temperature. A cocktail of primary antibodies, diluted in 0.5% BSA, was applied overnight at 4°C at the dilutions indicated in **Table 2**. The following day, slides were first washed with 0.05% PBS-Tween and then with PBS, followed by incubation with Iridium-conjugated intercalator (Fluidigm, catalogue number 201192B), diluted 1:2000 in 0.5% BSA for 30 minutes at room temperature. Lastly, slides were dipped in water (Invitrogen ultrapure distilled water), air dried and stored at room temperature until they were ablated.

#### Identification of region of interest (ROI) for laser ablation

Two serial sections each stained for either IF or IMC, were used. Based on IF staining with an antibody specific for proteolipid protein (PLP) (to visualize myelin) and DAPI (to visualize nuclei), ROIs were selected for ablation to capture the regions of interest for this study.

#### High-spatial resolution laser ablation of tissue sections

Tissue sections were analyzed by IMC, which couples laser ablation techniques and CyTOF mass spectrometry[2] (Cytof software v6.7). Briefly, a UV laser beam (λ = 219 nm) with a 1µmx1µm spot size is used to ablate the tissue. The laser rasters across the tissue at a rate of 200Hz (200 pixels/s) with the requisitie energy to fully remove the tissue within the selected region of interest. The ablated tissue is then carried by a stream of inert helium and argon gas into the Helios (a CyTOF system) where the material is completely ionized in the inductively coupled plasma. The ionized material then passes through high pass ion optics to remove ions with a mass less than 75amu before the ions enter the time of flight detector where they are separated based on their mass[6,4].

#### Data analysis and image visualization

Images of each mass channel were reconstructed by plotting the laser shot signals in the same order they were recorded, line scan by line scan, generating pseudo-colored intensity maps of each mass channel. These data were examined using MCD Viewer (V.1.0.560, Fluidigm). For qualitative assessments, images remained at the automatic threshold generated by MCD Viewer, based on the on the 98th percentile of signal. For further analysis, data was exported from MCD Viewer as tiff files, and each channel was run through an individual analysis pipeline in CellProfiler[8,22] (V3.185) in order to despeckle the image. Composite images were created for each ROI using Image J (V1.52a), and any changes to the brightness or contrast of a given marker were consistent across ROIs.

#### Calculation of the limit of detection

MCD Viewer was used to export text files acquired with the Hyperion IMC instrument (Fluidigm Inc., Markham ON), which were then converted to 32-bit single-channel TIFF images. The polygon tool within ImageJ 1.15s was used to manually outline the ROI (white matter of control, normal-appearing white matter) or subROI (periplaque white matter, lesion edge, lesion core), manually identified on the bases of PLP, HLA and Iridium-intercalator signals. Grey matter was excluded from subsequent analysis. Each image was despeckled in Definiens Developer XD 2.7 (Definiens Inc, Munich, Germany), using a 2D gray-level morphological opening filter with kernel radius of 1. In addition, the intensities of each marker were normalized using a modified z score approach, in which the intensity of each pixel is divided by the sum of (mean intensity of the image plus 3 times the standard deviation of the pixels in the image) I_zs_ = I/(μ_Im_+3\**σ*_Im_). This normalization approach has been previously used [13] and we found that it allows for a reliable comparison between IMC markers across different channels, with per-marker comparisons holding robustly across a 16-fold antibody dilution series (data not shown).

#### Single-cell segmentation

In order to define cells, we used a customized segmentation algorithm that took into account both the presence of nuclear DNA Iridium-intercalator as well as a set of markers of interest (see example in **Figure 7 – Supplement Figure 1**). In brief, a Gaussian blur was applied to the DNA signal and the resulting blurred image was segmented to identify nuclear content (**Figure 7 – Supplement Figure 1a**). Segmentation around the nuclei was expanded to simulate the cytoplasm, corresponding to individual cell areas, using a combination of threshold and watershed filters (**Figure 7 – Supplement Figure 1b**). Next we interrogated the segmented image for the presence of specific markers or combinations of markers that are either biologically co-expressed, or whose expression is mutually exclusive, according to the combination of markers indicated in **Table 3**. If a nucleated cell was positive for a marker or a combination of markers (see example for CD3 in **Figure 7 – Supplement Figure 1c**), the marker(s) signal was used to refine the initial nuclear segmentation. Nucleated cells that were not positive for any of the markers used, were segmented purely based on DNA signal and expanded to simulate the cytoplasmic area around the nucleus.

**Table 3.** Antibody panels for the identification of functional cell types by IMC.

| Functional Cell Type | Antibody Panel |
| --- | --- |
| **Macrophages and microglia** |  |
| Macrophages | CD45<sup>high</sup>HLA<sup>+</sup>TMEM119<sup>-</sup> |
| Activated macrophages | CD45<sup>high</sup>HLA<sup>+</sup>TMEM119<sup>-</sup>CD68<sup>+</sup>PLP<sup>-</sup> |
| Demyelinating macrophages | CD45<sup>high</sup>HLA<sup>+</sup>TMEM119<sup>-</sup>CD68<sup>+</sup>PLP<sup>+</sup> |
| Microglia | CD45<sup>low/+</sup>HLA<sup>+</sup>TMEM119<sup>+</sup> |
| Activated microglia | CD45<sup>low/+</sup>HLA<sup>+</sup>TMEM119<sup>+</sup>CD68<sup>+</sup>PLP<sup>-</sup> |
| Demyelinating microglia | CD45<sup>low/+</sup>HLA<sup>+</sup>TMEM119<sup>+</sup>CD68<sup>+</sup>PLP<sup>+</sup> |
| **T Cells** |  |
| CD8<sup>+</sup> T cells | CD45<sup>+</sup>CD3<sup>+</sup>CD8α<sup>+</sup>CD4<sup>-</sup> |
| Proliferating CD8<sup>+</sup> T cells | CD45<sup>+</sup>CD8α<sup>+</sup>CD3<sup>+</sup>CD4<sup>-</sup>Ki67<sup>+</sup> |
| Activated CD8<sup>+</sup> T cells | CD45<sup>+</sup>CD3<sup>+</sup>CD8α<sup>+</sup>CD4<sup>-</sup>NFAT<sup>+</sup> |
| Activated and proliferating CD8<sup>+</sup> T cells | CD45<sup>+</sup>CD3<sup>+</sup>CD8α<sup>+</sup>CD4<sup>-</sup>NFAT<sup>+</sup>Ki67<sup>+</sup> |
| “Chronically activated” CD8<sup>+</sup> T cells | CD45<sup>+</sup>CD3<sup>+</sup>CD8α<sup>-</sup>CD8<sup>+</sup>CD38<sup>+</sup>HLA<sup>+</sup> |
| CD4<sup>+</sup> T cells | CD45<sup>+</sup>CD3<sup>+</sup>CD8α<sup>-</sup>CD4<sup>+</sup> |
| Proliferating CD4<sup>+</sup> T cells | CD45<sup>+</sup>CD8α<sup>-</sup>CD3<sup>+</sup>CD4<sup>+</sup>Ki67<sup>+</sup> |
| Activated CD4<sup>+</sup> T cells | CD45<sup>+</sup>CD3<sup>+</sup>CD8α<sup>-</sup>CD4<sup>+</sup>NFAT<sup>+</sup> |
| Activated and proliferating CD4<sup>+</sup> T cells | CD45<sup>+</sup>CD3<sup>+</sup>CD8α<sup>-</sup>CD4<sup>+</sup>NFAT<sup>+</sup>Ki67<sup>+</sup> |
| “Chronically activated” CD4<sup>+</sup> T cells | CD45<sup>+</sup>CD3<sup>+</sup>CD8α<sup>-</sup>CD4<sup>+</sup>CD38<sup>+</sup>HLA<sup>+</sup> |
| **B Cells** |  |
| Naïve B Cells | CD45<sup>+</sup>Igκ/Igλ<sup>+</sup>IgM<sup>+</sup>CD38<sup>-</sup> |
| IgM memory B cells | CD45<sup>+</sup>Igκ/Igλ<sup>+</sup>IgM<sup>+</sup>CD38<sup>+</sup> |
| Switched memory B cells | CD45<sup>+</sup>Igκ/Igλ<sup>+</sup>IgM<sup>-</sup>CD38<sup>+</sup> |

#### Gating strategy for quantitative analysis of T cell, B cell, macrophage and microglial cell subsets

A segmented cell export of raw and normalized marker intensities for all channels in each region of interest were exported as a single csv file. The per-cell mean intensities of each marker combination, (see marker list in **Table 3**), were linearly rescaled for visualization purposes. 2D log-log biaxial scatterplots of these intensities were generated in Python (V3.6.8) using matplotlib (V3.0.3). A positive- and negative-gating strategy was applied to establish thresholds that identify particular cell types (see **Figure 7 – Supplement Figure 2** and the method below). Quadrants were set on pathologist-verified positive cells. In brief, ROI were examined in Definiens Developer XD 2.7. Cells were manually annotated by a pathologist, based on the expression of a biologically relevant set of markers to identify cells in each class of interest as defined below. These identified positive cells were superimposed to the 2D log-log scatterplots to definitively establish gates that would capture the appropriate positivity range of each cell phenotype as shown in **Figure 7 – Supplement Figure 2**.

#### For T cells

All nucleated cells expressing Igκ, Igλ, IgM, CD68 and HLA were eliminated, as these markers are not expressed on T cells. Gates were established for CD3 and CD45 based on a 2D log-log scatterplot of these markers. Following the identification of T-lineage cells, the same procedure was performed for CD3 vs CD4 and CD3 vs CD8, resulting in the identification of two subpopulations: CD4^+^ T cells and CD8^+^ T cells. Thresholds for Ki67 (a marker of proliferation) and NFAT1 (a marker of activation) were established based on manually annotated CD3^+^KI67^+^ and CD3^+^NFAT1^+^ cells, as described above. All cell populations were validated by manual annotation as described above.

#### For B Cells

All nucleated cells expressing CD3, CD4, CD8 and CD68 were eliminated as these markers are not expressed on B cells. B cells were further identified by CD45 above the same threshold set for T cells. Scatterplot comparison for Igκ and Igλ intensity identified Igκ^+^ and Igλ^+^ single-positive populations. Igκ^+^Igλ^+^ double positive cells were eliminated as artifactual, since the two allelic variant cannot co-exist on a given cell. Within Igκ^+^ or Igλ^+^ B-lineage cells, we compared IgM to CD38 to determine the relative abundance of IgM^+^ or CD38^+^ cell subpopulations. All cell populations were validated by manual annotation as described above.

#### For macrophages and microglia

All nucleated cells expressing CD3, CD4, CD8, Igκ/λ and IgM were eliminated as these markers are not expressed on macrophages and microglia. Discrimination of the remaining cells was visualized in a scatterplot for TMEM119 and CD45. The threshold for TMEM119 positive signal was determined by comparison to TMEM119^+^ microglial cells that were identified by manual observation, relative to other cell types. The threshold for CD45 high or low signal was determined by the comparison to manually identified TMEM^−^ macrophages. Manually identified microglial cells were used to establish the lower limit of the CD45 quadrants. Cells that were low for both TMEM119 and CD45 were labeled “other” and ignored from subsequent analysis. These latter cells, likely correspond to astrocytes, oligodendrocytes and other cell types. Both TMEM119^+^CD45^low/+^ microglial cells and TMEM^−^CD45^high^ macrophages were further evaluated for HLA, CD68 and PLP (depicted as scatterplots), to differentiate microglia or macrophages that are either resting or activated/phagocytic/demyelinating. All cell populationes were validated by manual annotation as described above.

#### Generation of cell density map

The gating strategy described above was confirmed by plotting the appropriately gated cell types for major lineage markers (see examples in **Figure 7 – Supplement Figure 3**). Note that Igκ > Igλ in **Figure 7 – Supplement Figure 3a** consistent with over-representatino of κ^+^ B cells in humans[24]. Following this confirmation, the density of all relevant cell subtypes was computed within each biological region of interest. A heat map, generated using Seaborn (V0.9.0), displayed the cell counts per mm^2^ of tissue.

#### Generation of distance map

To assess the location of identified cells relative to blood vessels, collagen^+^ perivascular regions of >10µm diameter and >800 µm^2^ area were segmented, and the distance between cells of interest and the border of these perivascular regions was calculated. Similarly to the cellular density calculations, average vessel distances corresponding to the mean of the per-cell vessel distance values were computed, expressed in µm, and presented as a distance heat map. Some regions did not contain any cells of a particular type, leading to undefined values for those particular regions and cell type combinations (presented as “n/a”, not applicable).

### Statistical analysis

Where statistical testing was possible, all tests were performed using Prism software (v5.01; GraphPadSoftware, San Diego, CA). Data distribution was tested for normality. Because all variables were not normally distributed, possible correlations between the density of nuclei or the density of CD3^+^ cells (number of cells / mm^2^ of tissue) detected either by IF or by IMC, were investigated with the nonparametric Spearman rank correlation. Differences were considered significant at p < 0.05.

## RESULTS

### Comparability of IF versus IMC approach and specificity of metal-conjugated antibodies on brain-resident cell types

In this study, we performed two separate IMC runs on the same patient and control tissue. The first run (Figure 1–2) was to validate our panel and the approach in general. The second run (Figure 3–7) was to evaluate the composition of immune cells across several types of lesions and sub-lesional areas. To evaluate the validity of IMC for the analysis of post-mortem MS brain tissue, we first investigated whether images generated by IMC revealed a similar number of cells expressing a given marker in a mm^2^ area of tissue, as determined by IF. We used IF or IMC on serial sections from the same tissue block. Sections were stained with DAPI (IF) or Iridium-intercalator (IMC) to identify DNA in the nuclei, and with anti-CD3-FITC (IF) or anti-CD3-170Er (IMC) to identify CD3^+^ T cells. Imaging of equivalent ROI in IF-and IMC-stained sections showed similar staining patterns with clearly resolved anatomical regions (**Figure 1a**). We also showed that IMC is able to resolve myelin engulfed by microglia/macrophages with a similar pattern as what is observed using IHC and IF approaches (**Figure 1 – Supplement Figure 3**). Quantification analysis showed that the number of nuclei or CD3^+^ T cells detected by IMC is on average 1.4 and 1.5 fold higher than numbers detected by IF, respectively. The increase in detection of cell counts across ROI for DNA and for CD3 using the IMC method compared to IF is either due to the necessary use of a serial section for comparison between the two techniques, or alternatively IMC may be a more sensitive approach for detecting nucleated cells within brain tissue. We also compared the number of nuclei identified with DAPI by IF and the number of nuclei identified with intercalator by IMC, as well as the number of CD3^+^ T cells identified with FITC-labeled secondary antibody by IF and the number of CD3^+^ T cells identified with the 170Er metal-tag by IMC. Both comparisons revealed a significant positive correlation (Spearman correlation coefficient: r=0.9182, p=0.0002 and r=0.8929, p=0.01, for nuclei and for CD3^+^ T cells respectively) (**Figure 1b, c**), indicating proportional representation of brain-resident cells was in agreement for both methods. Collectively, IMC reproduces staining patterns that are in agreement with those produced using a standard IF method in MS brain tissue.

**Figure 1.**
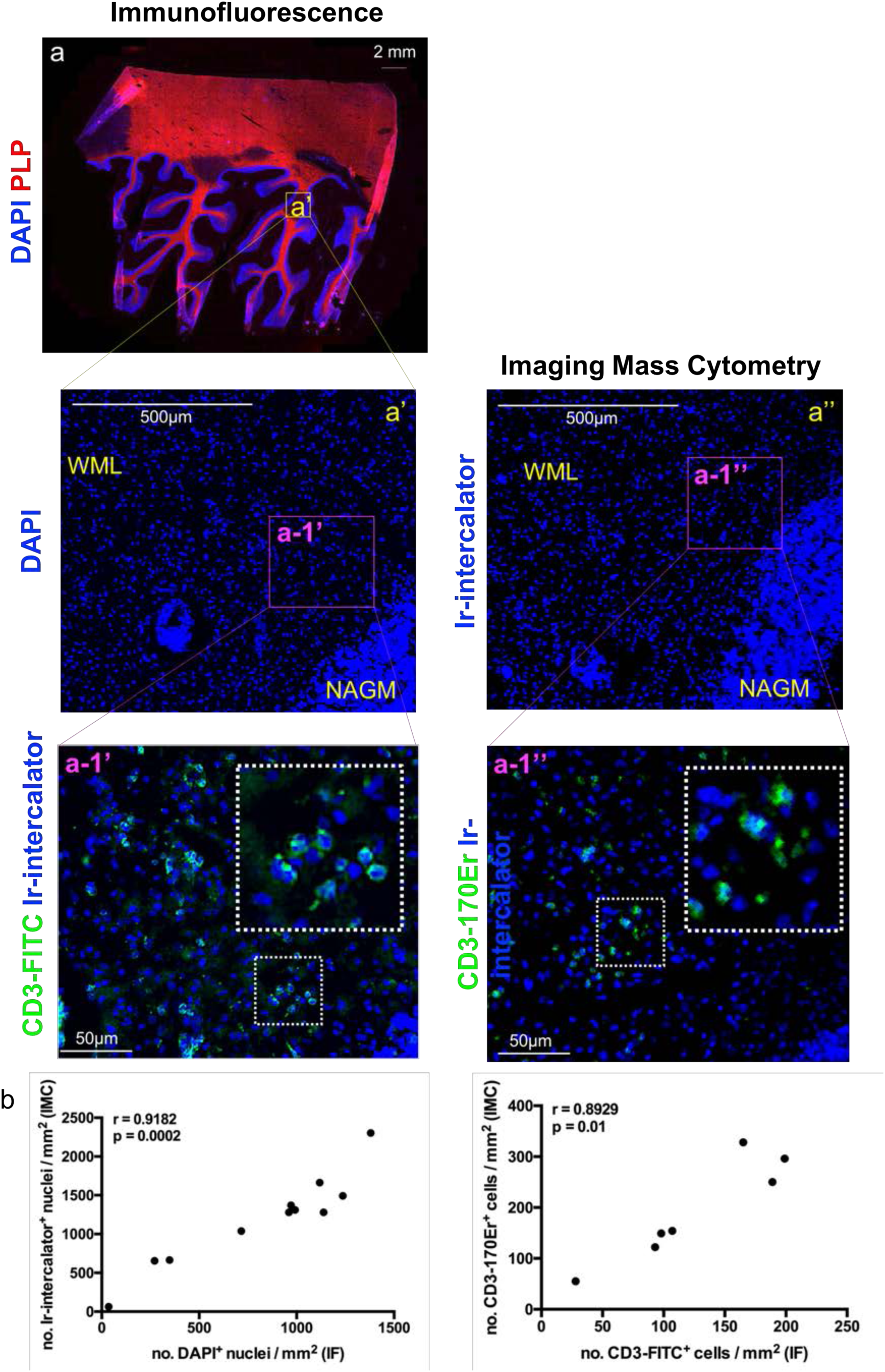
Comparing IMC to IF in MS lesions. Two serial sections were assessed: one used for immunofluorescence (IF, **a and a’**) and one dedicated to Imaging Mass Cytometry (IMC, **a”**). The region of interest (**a’**) was guided by the immunofluorescence staining with anti-PLP (proteolipid protein, shown in red to visualize myelin), and DAPI (shown in blue to visualize nuclei) (**a**) for the identification of lesion location and type (see **Figure 1 – Figure Supplement 1**). The entire region of interest on a serial section was subjected to IMC, according to the work flow shown in **Figure 1 – Figure Supplement 2**. Staining with Iridium (Ir)-intercalator is shown in blue to visualize DNA in nuclei. A blow up area (referred to as **a-1’** for IF and **a-1”** for IMC) of the region of interest within an active lesion, was also stained with fluorochrome conjugated anti-CD3 (a-1’) or metal conjugated anti-CD3 (a-1”), both depicted in green. WML, white matter lesion; NAGM, normal-appearing grey matter. (**b**) Spearman correlation coefficient, showing a significant positive correlation between the number of nuclei identified with DAPI by IF and the number of nuclei identified with Ir-intercalator by IMC (n=11, coefficient, r=0.9182, p=0.0002). (**c**) Spearman correlation coefficient, showing a significant positive correlation between the number of CD3^+^ T cells identified with fluorochrome-conjugated antibody by IF and the number of CD3^+^ T cells identified with metal-conjugated antibody by IMC (n=7, coefficient, r=0.8929, p=0.01).

**Figure 2.**
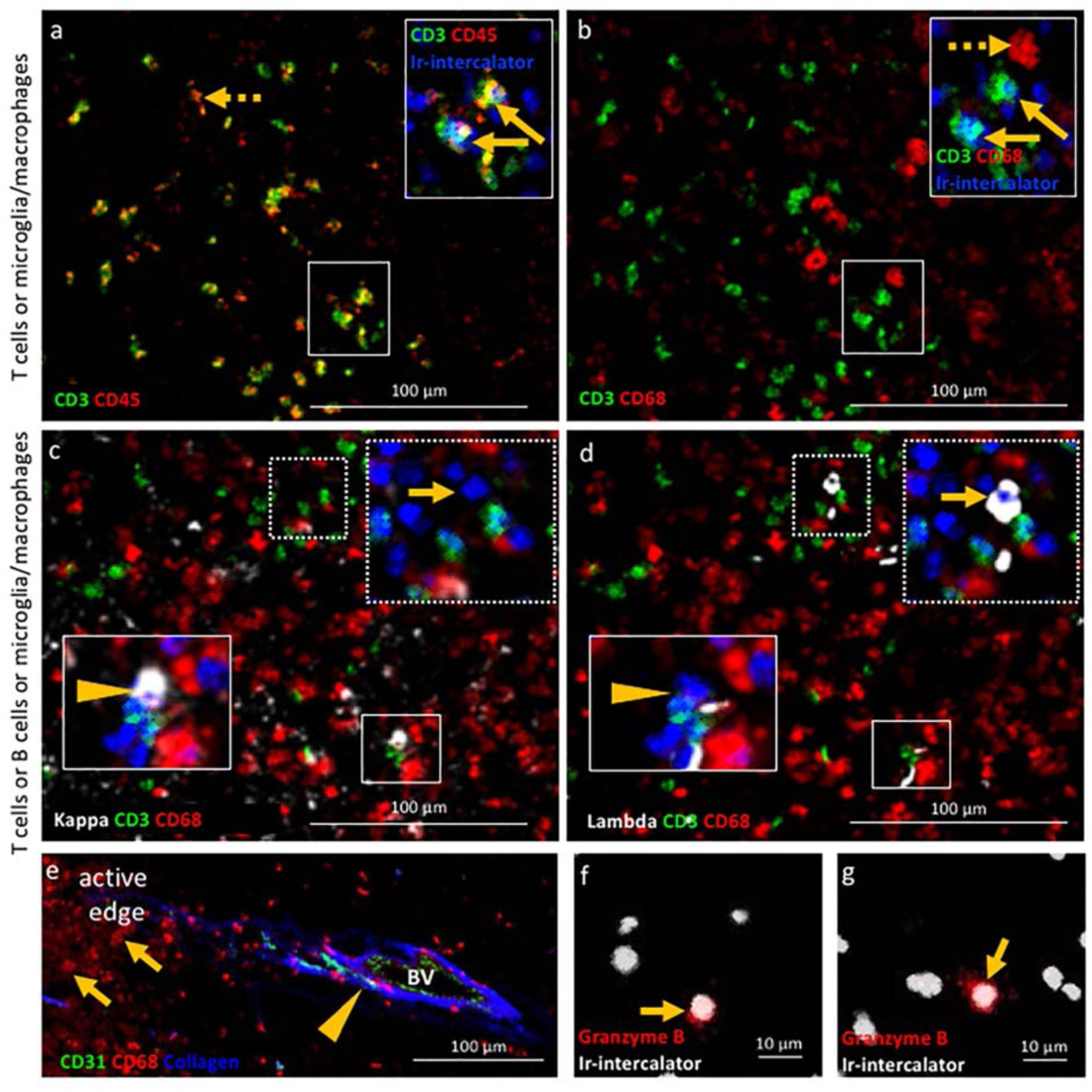
Validation of IMC specificity in MS lesions. (**a**) Overlay of CD3 (green) and CD45 (red) identifies CD3^+^CD45^+^ T cells (solid arrows) and CD3^−^CD45^+^ leukocytes other than T cells (dotted arrow). (**b**) Overlay of CD3 (green) and CD68 (red) identifies CD3^+^CD68^−^ T cells (solid arrows) and CD3^−^CD68^+^ microglia /macrophages (dotted arrow). Note that the solid arrows in a and b indicates the same CD3^+^CD45^+^CD68^−^ T cells. (**c**) Overlay of κ (white), CD3 (green) and CD68 (red) and (**d**) overlay of λ (white), CD3 (green) and CD68 (red) identify κ^+^CD3^−^CD68^−^ B cells (arrow head in **c**) that are λ^−^CD3^−^CD68^−^ (arrow head in **d**) and κ^−^CD3^−^ CD68^−^ B cells (arrow in **c**) that are λ^+^CD3^−^CD68^−^ (arrow in **d**), as expected based on the allelic exclusion of κ and λ. (**e**) Overlay of CD31 (green), CD68 (red) and Collagen (blue) identifies CD31^+^Collagen^+^CD68^−^ endothelial cells (arrow head) and CD31^−^Collagen^−^ CD68^+^ microglia/macrophages (arrows). (**f, g**) Granzyme B^+^ cells (arrows). Images in **a** and **b** as well as images in **c** and **d** are from the same areas of an active demyelinating lesion. Image in **e** are from the edge of an active demyelinating lesion. Images in **f** and **g** are from the center of an active demyelinating lesion.

**Figure 3.**
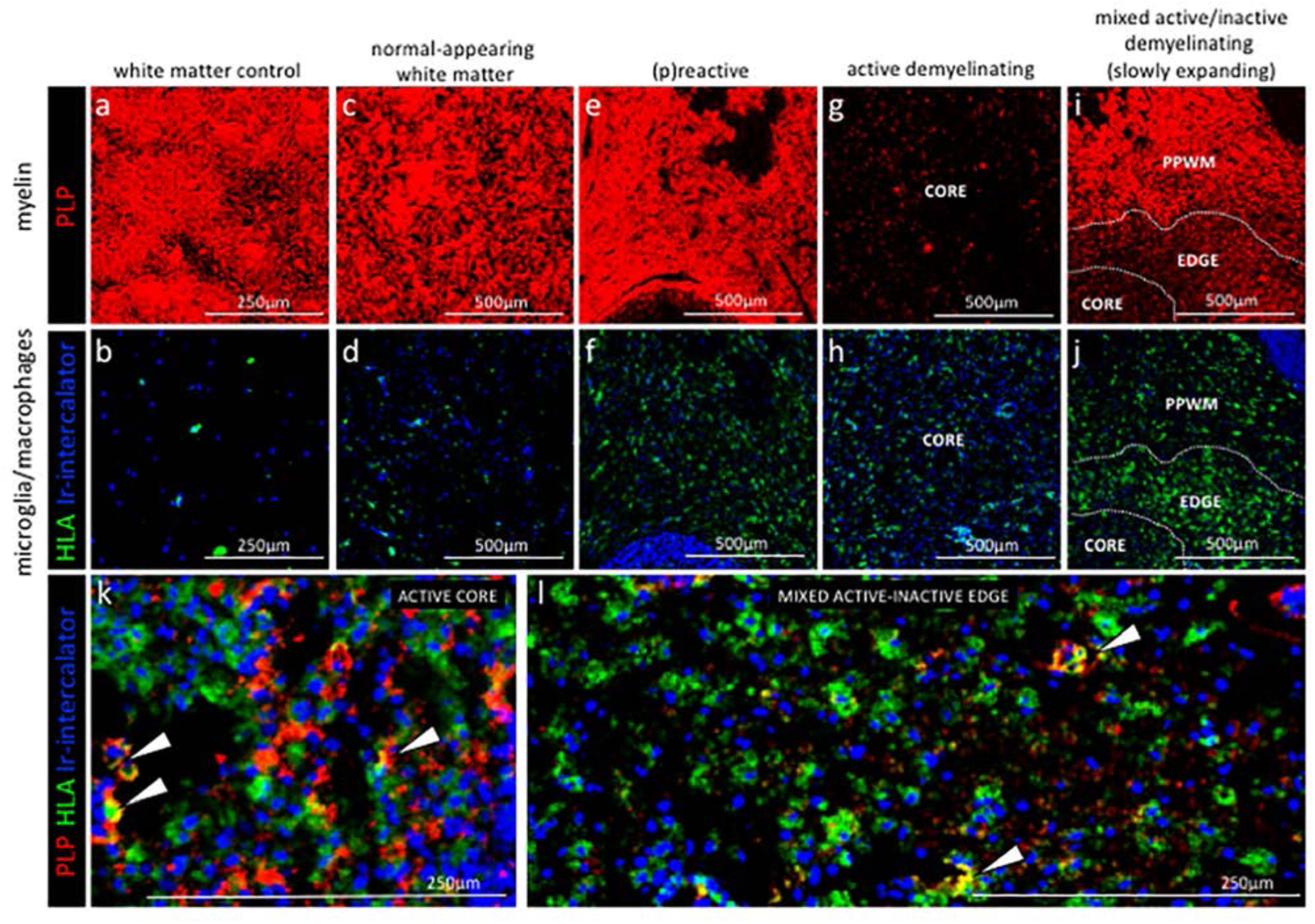
Staging of MS lesions by IMC. Representative mass cytometry images of white matter areas of healthy control (**a, f**), MS normal-appearing white matter (block no. CR4A) (**b, g**), MS (p)reactive lesion (block no. CR4A) (**c, h**), MS active demyelinating lesion (block no. CR4A) (**d-i**) and an MS slowly expanding lesion (block no. CL3A) (**e-j**). For each region of interest, we show the same area simultaneously labeled with markers of myelin (proteolipid protein, PLP), antigen presentation (human leukocyte antigen, HLA) to detect microglia/macrophages and DNA (intercalator). Images of PLP (red) (**a-e**) and overlay of HLA (green) and intercalator (blue) (**f-i**) show the lesion activity in staged MS lesions compared to control white matter and normal-appearing white matter. (**k, l**) Overlay of PLP, HLA and intercalator show microglia/macrophages containing PLP^+^ myelin protein in the core of an active lesion (**k**) and in the edge of a slowly expanding lesion (**l**), indicative of demyelinating activity. PPWM, periplaque white matter; BV, blood vessel.

Second we assessed the target specificity of metal-tagged antibodies by IMC. We stained brain tissue with metal-tagged antibodies against molecules that are either expected to be co-expressed by cells or whose cellular expression is expected to be mutually exclusive. IMC imaging of the edge of an active demyelinating lesion identified CD3^+^CD45^+^ T cells and CD3^−^ CD45^+^ leukocytes other than T cells (**Figure 2a**). The co-expression of CD68 in the latter cell types identifies these as microglia/macrophages (CD3^−^CD45^+^CD68^+^) (**Figure 2b**). As expected, CD3^+^ T cells lack expression of immunoglobulin light chain, (CD3^+^κ/λ^−^). Since antibodies directed to the B cell-restricted lineage markers CD19 and CD20 were sub-optimal on our brain tissues, we relied instead on antibodies that detect the two allelic variants of the immunoglobulin light chain (*κ/λ*). This provided two advantages: the ability to capture all B cells, irrespective of their maturation/activation status (for example, plasma cells downregulate CD19/CD20) and the ability to test specificity of our B cell directed reagents since κ and λ are allelically excluded on the surface of B cells. As expected, we identified B cells that were either positive for the κ or the λ light chain of immunoglobulins and negative for the CD3 marker of T cells and the CD68 marker of macrophages (for example, CD3^−^CD68^−^κ^+^ B cells in **Figure 2c arrow**, versus CD3^−^CD68^−^λ^−^ B cells in **Figure 2d arrow** and vice versa CD3^−^CD68^−^κ^−^ B cells in **Figure 2c arrow head**, versus CD3^−^CD68^−^λ^+^ B cells in **Figure 2d arrow head**). We found that the extracellular matrix protein collagen (not cell-associated) surrounded putative blood vessels lined by endothelial cells that expressed the CD31 marker as expected. In contrast, macrophages visualized by CD68 do not stain positive for collagen nor express CD31 (CD31^−^ collagen^−^CD68^+^) (**Figure 2e**). Lastly, we asked whether the IMC approach would have sufficient sensitivity to detect soluble molecules that can be rare in tissues. For this, we used an antibody against granzyme B, a serine protease with pro-inflammatory function produced by activated cytotoxic T cells. IMC identified a granzyme B^+^ signal with granular expression in close proximity to nuclei (**Figure 2f, g**). These results show that IMC enables imaging of multiple markers on a single tissue section reproducing IF-equivalent staining patterns and with cell lineage-specific markers expressed on appropriate cell types.

### Qualitative staging of MS lesions by IMC

To verify whether IMC provides us with the ability to differentiate normal-appearing tissue versus different lesional stages of the MS brain, we analysed the proteolipid protein (PLP) signal that visualizes myelin and the human leukocyte antigen (HLA) signal that visualizes antigen presenting cells (**Figure 3**). Note that in some ROI, the PLP staining pattern reflects the cross-sectional orientation of myelinated fibers (for example WMC and NAWM), whereas in others, longitudinal myelin tracks are observed (for example (p)reactive). The different orientation of the tissue results from the sectioning plane of the tissue block and is reflected in the staining pattern displayed. Consistent with the generally non-inflamed and myelinated state of healthy white matter, control white matter showed intact myelin staining with few HLA^+^ cells (**Figure 3 a, b**). Normal appearing white matter (NAWM) in the MS brain exhibited normal myelin staining, however HLA^+^ cells within the MS NAWM appeared enriched when compared to control white matter (**Figure 3 c, d**). Similarly, the (p)reactive lesion showed a normal myelin signal but HLA^+^ cells accumulated at this site (**Figure 3 e, f**). The active lesion core showed loss of PLP signal with accumulation of HLA^+^ cells (**Figure 3 g, h**). The mixed active-inactive lesions showed reduced myelin and accumulation of HLA^+^ cells at the lesion edge (**Figure 3 i, j**). HLA^+^ cells that contained PLP myelin products were found in both active lesions[26] (**Figure 3k**) and at the edge of mixed active-inactive lesions (**Figure 3l**), indicative of demyelinating activity. Collectively we were able to show that IMC of different ROI (pre-selected on the bases of PLP/HLA IF staining on a serial section) was able to differentiate between normal-appearing tissue and different lesional stages of the MS brain.

### Qualitative assessment of microglia and macrophages in staged MS lesions by IMC

Next, we analysed key molecules that differentiate between the phenotype and functional status of microglia and macrophages in relation to the lesional stage and demyelinating activity of MS lesions.

#### Control subject white matter

In the white matter from a control subject we found that microglia, identified as being TMEM119^+^, generally showed a thin ramified morphology, typical of resting cells (**Figure 4 a, a’ dotted arrows**). On these cells, the HLA marker of antigen presentation was generally low or not detectable, confirming a quiescent state. On the contrary, TMEM119^+^ microglia that showed a more rounded morphology, which is a sign of activation, also stained for HLA and CD68 (**Figure 4 a, a’ arrow head and b, b’ arrow head**). HLA/CD68 expression is indicative of antigen presentation and phagocytic activity, respectively. TMEM119^−^HLA^+^CD68^+^ cells were identified as macrophages and were also present in the white matter of control (**Figure 4 a, a’ arrow and b, b’ arrow**). These data indicate that in the normal white matter of a control subject some microglia (TMEM119^+^) and some macrophages (TMEM119^−^) have an activated phenotype (HLA^+^CD68^+^).

**Figure 4.**
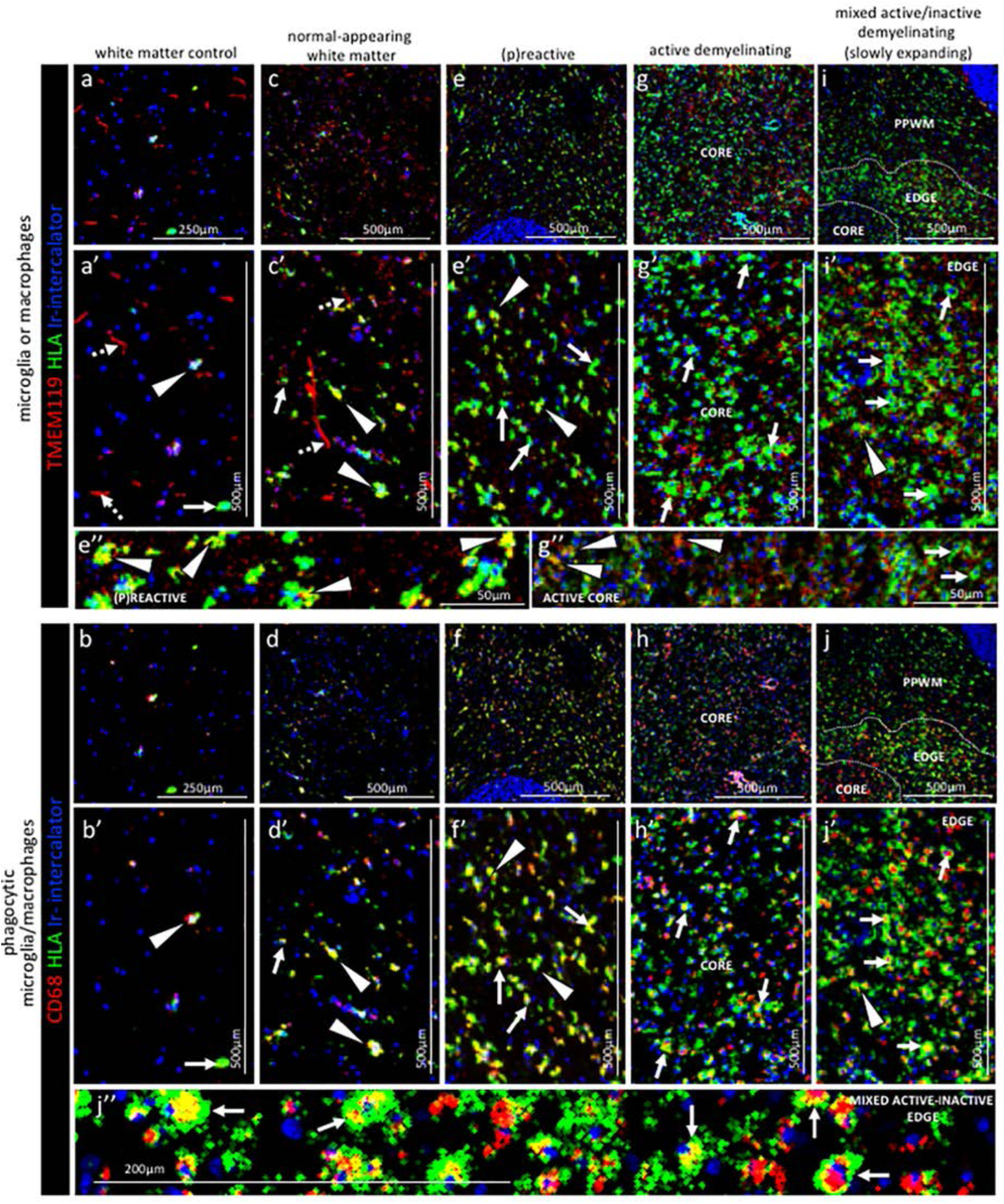
Pattern of microglia or macrophage activity in different stages of MS lesions by IMC. Representative mass cytometry images of control white matter (**a, a’, b, b’**), normal-appearing white matter (block no. CR4A) (**c, c’, d, d’**), (p)reactive lesion (block no. CR4A) (**e, e’, f, f’**), active demyelinating lesion (block no. CR4A) (**g, g’, h, h’**) and slowly expanding lesion (block no. CL3A) (i**, i’, j, j’**). For each region of interest, we show the same area simultaneously labeled with markers of antigen presentation (human leukocyte antigen, HLA) to detect microglia and/or macrophages, TMEM119 to detect microglia, lysosomes (CD68) to detect phagocytic cells and DNA (Ir-intercalator). (**a, a’-i, i’**) Overlay of TMEM119 (red), HLA (green) and Ir-intercalator (blue) identifies TMEM119^+^HLA^−^ resting microglia with thin elongated processes (dotted arrows in **a’ and c’**) and TMEM119^+^HLA^+^ activated microglia (arrows head in **a’,c’, e’, i’ and e”**) or TMEM119^−^HLA^+^ activated macrophages (solid arrows in **a’, c’, e’, g’, i’ and g”**). (**b, b’-j, j”**) Overlay of CD68 (red), HLA (green) and intercalator (blue) identifies HLA^+^CD68^+^ phagocytic microglia/macrophages. PPWM, periplaque white matter; BV, blood vessel.

#### Normal-appearing white matter

Visualization of the expression pattern of microglia and macrophage markers in the normal-appearing white matter showed some TMEM119^+^ microglia with ramified morphology (**Figure 4 c, c’ dotted arrows**), similar to control white matter. However unlike control white matter, the normal-appearing white matter showed many TMEM119^+^ microglia that were also positive for HLA and CD68 (**Figure 4 c, c’ arrows head and d, d’ arrows head**). A few TMEM119^−^HLA^+^CD68^+^ macrophages were also present in the normal-appearing tissue (**Figure 4 c, c’ arrow and d, d’ arrow**).

#### (P)reactive lesions

Within the (p)reactive lesions, TMEM119^+^ microglia accumulated, showed an enlarged morphology that is indicative of an activated state, and expressed both HLA and CD68 (**Figure 4 e, e’ and e” arrows head and f, f’ arrows head**). TMEM119^−^ HLA^+^CD68^+^ macrophages were also present (**Figure 4 e, e’ arrows and f, f’ arrows**).

#### Active lesions

Active lesions contained high numbers of TMEM119^+^HLA^+^CD68^+^ microglia and TMEM119^−^HLA^+^CD68^+^ macrophages, most of them with enlarged and foamy morphology that is typical of the activated and phagocytic state (**Figure 4 g, g’ and g” arrows and h, h’ arrows**).

#### Mixed active-inactive lesion (slowly expanding lesion)

The edge of these lesions was characterized by a rim of dense TMEM119^+^HLA^+^ microglia (**Figure 4 i, i’ arrow head and j, j’ arrow head**) and TMEM119^−^HLA^+^ macrophages (**Figure 4 i, i’ arrows and j, j’, j” arrows**), both with obvious enlarged CD68^+^ lysosomes (**Figure 4 j” arrows**). Only a few HLA^+^CD68^+^ cells were present in the inactive lesion core and microglia showed profound reduction in the HLA signal (**Figure 4 i, i’ arrows and j, j’, j” arrows**).

### Qualitative assessment of T cells in staged MS lesions by IMC

Next, we analysed key molecules that differentiate between the phenotype and functional status of T cells in relation to the lesional stage and demyelinating activity of MS lesions.

#### Control subject white matter

In the white matter from a control subject CD3^+^CD8α^−^ T cells or CD3^+^CD8α^+^ T cells were rare of absent (**Figure 5 a, a’ and b, b’**).

**Figure 5.**
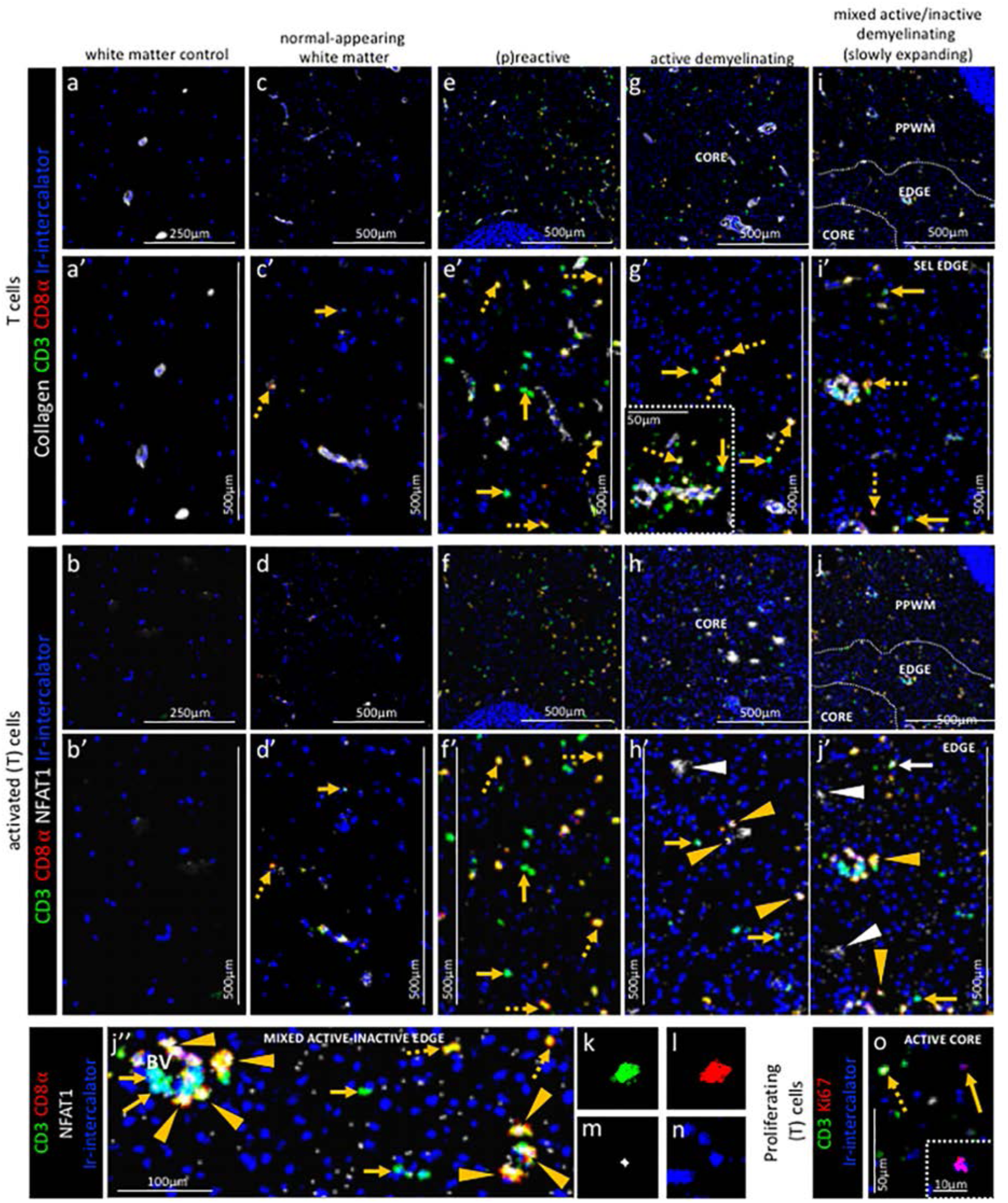
Pattern of T cell subpopulations in different stages of MS lesions by IMC. Representative mass cytometry images of white matter of control (**a, a’, b, b’**), normal-appearing white matter (block no. CR4A) (**c, c’, d, d’**), (p)reactive lesion (block no. CR4A) (**e, e’, f, f’**), active demyelinating lesion (block no. CR4A) (**g, g’, h, h’, o**) and slowly expanding lesion (block no. CL3A) (**i, i’, j-n**). For each region of interest, we show the same area simultaneously labeled with anti-collagen antibodies to visuzliae blood vessels, all T cells (CD3), CD8α T cells, cell proliferation (Ki67) and DNA (Ir-intercalator). (**a, a’-i, i’**) Overlay of collagen (white), CD3 (green), CD8α (red) and Ir-intercalator (blue) identifies CD3^+^CD8α^+^ T cells (dotted arrows in **c’, e’, g’ and i’**), CD3^+^CD8α^−^ (therefore by exclusion putative CD4^+^) T cells (solid arrows in **c’, e’, g’ and i’**) and collagen^+^ blood vessels. (**b-b’-j, j”**) Overlay of CD3 (in green), CD8α (red), NFAT1 (in white) and Ir-intercalator (in blue) identifies CD3^+^CD8α^+^NFAT1^+^ T cells (yellow arrow head in **h’, j’ and j”**) and CD3^+^CD8α^−^NFAT1^+^ (putative CD4^+^) T cells (white solid arrow in **j’**). CD3^−^CD8α^−^NFAT1^+^ cells are also detected (white arrow head in **h’ and j’**). (**o**) Overlay of CD3 (in green), Ki67 (red) and Ir-intercalator (in blue) identifies CD3^+^Ki67^+^ proliferating Tcells (dotted arrow) and CD3^−^Ki67^+^ proliferating cells other than T cells (solid arrows and inset). PPWM, periplaque white matter.

#### Normal-appearing white matter

In the normal-appearing white matter, we identified some CD3^+^CD8α^−^ T cells (**Figure 5 c, c’ arrow**) and some CD3^+^CD8α^+^ T cells (**Figure 5 c, c’ dotted arrow**) that did not show signs of activation as defined by the expression of NFAT1 which translocates to the nucleus of T cells upon T cell receptor activation[30] (**Figure 5 d, d’ arrow and dotted arrow**).

#### (P)reactive lesions

Within the (p)reactive lesions, CD3^+^CD8α^−^ and CD3^+^CD8α^+^ T cells were both prominent (**Figure 5 e, e’ arrow and dotted arrow, respectively**) but did not stain for NFAT1 and were therefore presumably not activated (**Figure 5 f, f’ arrow and dotted arrow, respectively**).

#### Active lesions

Active lesions contained both, CD3^+^CD8α^−^ and CD3^+^CD8α^+^ T cells (**Figure 5 g, g’ arrow and dotted arrow, respectively**), mostly located in the perivascular area (**Figure 5 g’ inset)**, but also scattered in the parenchyma. Some CD3^+^CD8α^+^ T cells were also activated based on the expression of NFAT1 (**Figure 5 h, h’ arrow head**).

#### Mixed active-inactive lesion (slowly expanding lesion)

Similar to the core of active lesions, the edge of the slowly expanding lesions contained both, CD3^+^CD8α^−^ and CD3^+^CD8α^+^ T cells (**Figure 5 i, i’ arrow and dotted arrow, respectively**). A few CD3^+^CD8α^−^ T cells and some CD3^+^CD8α^+^ T cells were also NFAT1^+^ (**Figure 5 j, j’ white arrow and yellow arrow head, respectively**). These were found both in the perivascular area and in the parenchyma (**Figure 5 j”)**. The staining pattern of NFAT1 was consistent with the nuclear localization of this transcription factor (**Figure 5 k-n)**. Nuclear NFAT1 signal was also observed on CD3^−^ cells, consistent with reports of its localization of cells other than T cells[30] (**Figure 5 h, h’ and j, j’ white arrows head)**. Occasionally, we observed Ki67^+^ proliferating cells (**Figure 5 o, arrow and inset)**, some of which were CD3^+^ T cells (**Figure 5 o, dotted arrow**). In the inactive lesion core, we observed both scattered CD3^+^CD8α^−^ T cells and CD3^+^CD8α^+^ T cells (**Figure 5 i**).

### Qualitative assessment of B cells in staged MS lesions by IMC

Next we analysed key molecules that differentiate between the phenotype and functional status of B cells in relation to the lesional stage and demyelinating activity of MS lesions.

#### Control subject white matter

IgM staining on cells can be indicative of either naive B lymphocytes or IgM memory B cells. Therefore we first analysed the tissue for the presence of cell-associated IgM staining signal. In control white matter, IgM was not found in association with cells in the parenchyma but was only found in association with blood vessels, identified by the collagen staining. Further analysis showed that the IgM signal in the perivascular space co-localizes with the immunoglobulin light chain Igκ**/**Igλ, indicating that this IgM^+^ Igκ**/**Igλ^+^signal represents either naive or IgM memory B cells, or alternatively cell-free immunoglobulins (**Figure 6 a, a’ and b, b’ arrow head**).

**Figure 6.**
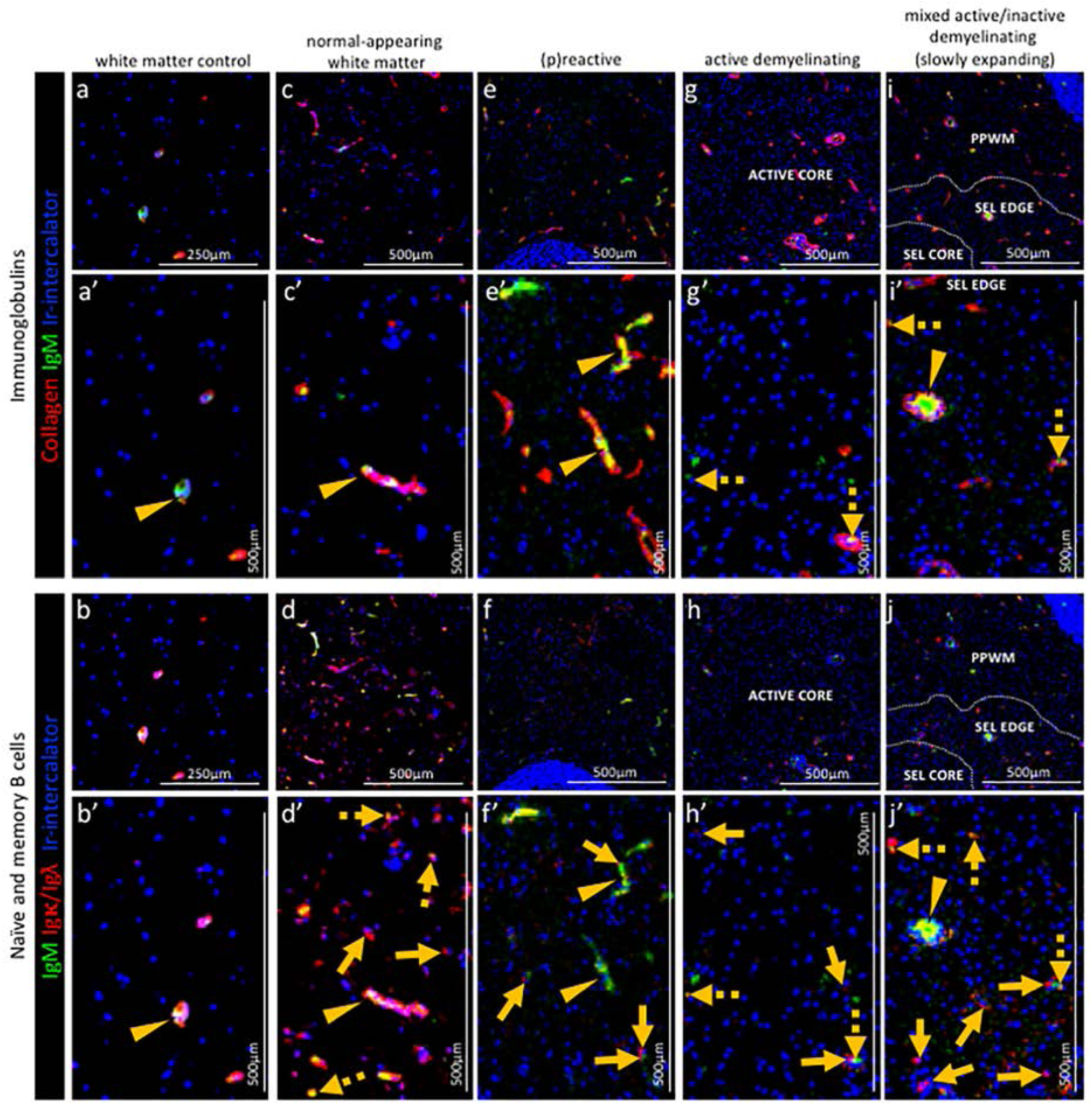
Pattern of immunoglobulins and B cell subpopulations in different stages of MS lesions by IMC. Representative mass cytometry images of white matter of control (**a, a’, b, b’**), normal-appearing white matter (block no. CR4A) (**c, c’, d, d’**), a (p)reactive lesion (block no. CR4A) (**e, e’, f, f’**), an active demyelinating lesion (block no. CR4A) (**g, g’, h, h’, o**) and a slowly expanding lesion (block no. CL3A) (i**, i’, j-n**). For each region of interest, we show the same area simultaneously labeled with markers of endothelial cells (collagen) to detect blood vessels, immunoglobulin M (IgM), the κ or λ light chain of immunoglobulins Igκ**/**Igλ to detect B cells and DNA (Ir-intercalator). (**a, a-i, i’**) Overlay of collagen (red), IgM (green) and Ir-intercalator (blue) identifies cellular (intercalator-associated, dotted arrows in g’ and i’) and non-cellular (free immunoglobulin, arrows head in **a’, c’, e’, i’**) IgM in the parenchyma or within collagen^+^ blood vessels. (**b, b’-j, j’**) Overlay of IgM (green), Igκ**/**Igλ (red) and Ir-intercalator (blue) identifies Igκ**/**Igλ^+^IgM^+^ naïve and IgM memory B cells (dotted arrow in **d’, h’ and j’**) and Igκ**/**Igλ^+^IgM^−^ class switch B cells (solid arrows in **d’, f’, h’ and j’**).

**Figure 7.**
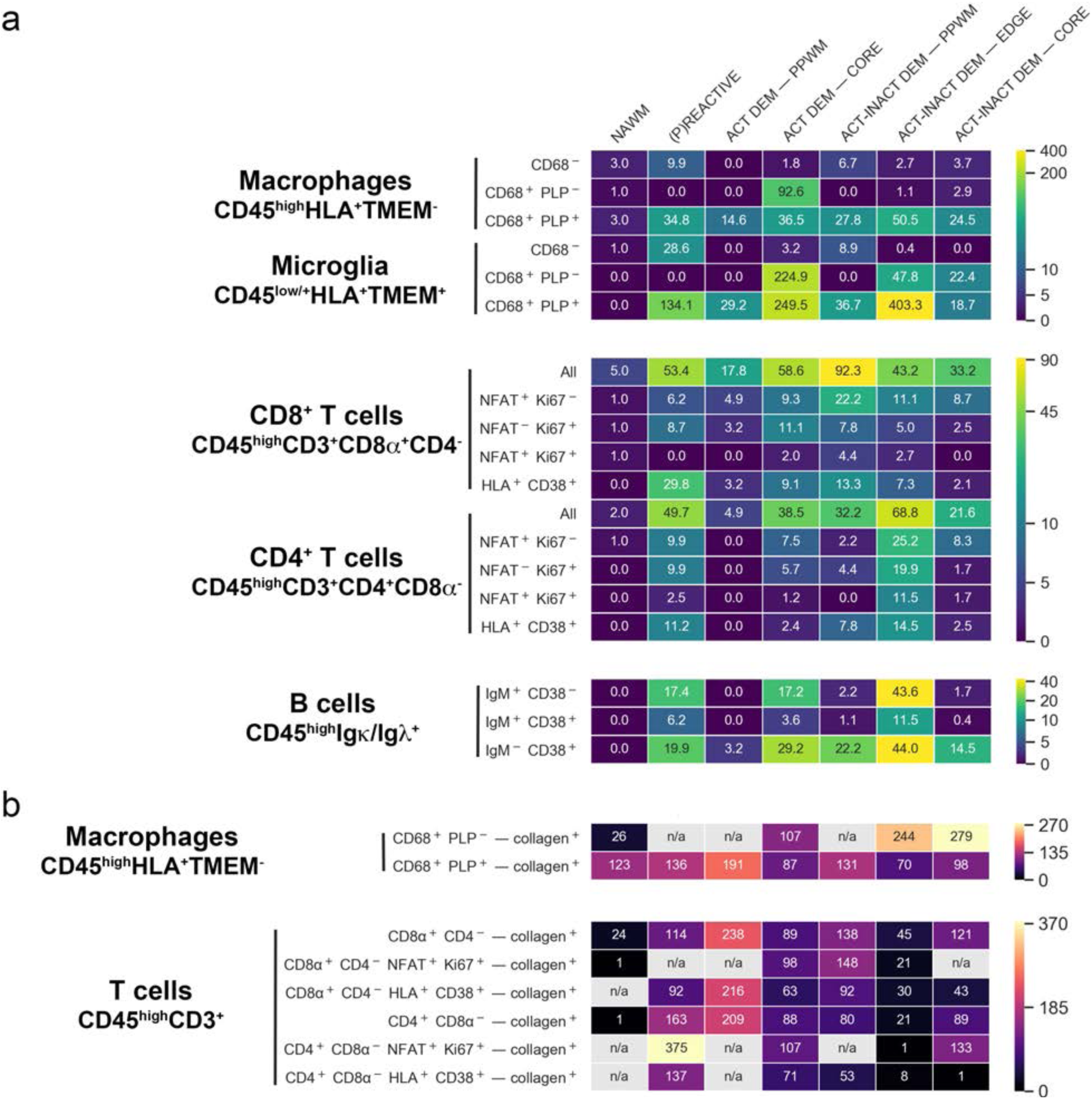
Density of immune cell subsets in different stages of MS lesions and their distance from blood vessels by IMC. (**a**) Cell counts are provided as number of cells per mm^2^ of region of interest. The category of cells is defined according to the expression of cell-specific and functional markers as indicated and also described in **Table 3**. (**b**) Distance between defined categories of cells and blood vessels (collagen^+^) are provided in μm. NAWM, normal-appearing white matter; PPWM, periplaque white matter; Act dem, active demyelinating; act inact dem, active-inactive demyelinating. The single-cell segmentation strategy is shown in **Figure 7 – Figure Supplement 1**. The Positive and negative “gates” used to identify each cell subset were established based on the quadrants laid out in **Figure 7 – Figure Supplement 2**. Please see the section “Gating strategy for quantitative analysis of T cell, B cell, macrophage and microglial cell subsets” in the materials and methods. The gating strategy used for the generation of heat maps is laid out in **Figure 7 – Figure Supplement 3**. Source files used for the quantitative analysis are provided in Figure 7 – Source data 1.

#### Normal-appearing white matter

In the normal-appearing white matter, the IgM signal was found both in association with blood vessels (**Figure 6 a, a’ and b, b’ arrow head**) and with nucleated Igκ**/**Igλ^+^ B cells scattered in the parenchyma (**Figure 6 a, a’ and b, b’ dotted arrows**). Nucleated Igκ**/**Igλ^+^ B cells that were IgM^−^ were also found in the parenchyma (**Figure 6 a, a’ and b, b’ solid arrows**) indicating the presence of class switched B cells.

#### (P)reactive lesions

Within the (p)reactive lesions, IgM was exclusively detected witin blood vessels (**Figure 6 e, e’ arrow head**). Nucleated Igκ**/**Igλ^+^ B cells were present and expressed a switched B cell phenotype (IgM^−^) (**Figure 6 e, e’ and f, f’ solid arrows**).

#### Active lesion

Within the active lesions, nucleated Igκ**/**Igλ^+^ B cells that displayed a switched phenotype (IgM^−^) were mostly present (**Figure 6 g, g’ and h, h’ solid arrows**).

#### Mixed active-inactive lesion (slowly expanding lesion)

Similarly to active lesions, at the edge of mixed active-inactive lesions, nucleated Igκ**/**Igλ^+^ B cells were present and displayed a switched memory phenotype (IgM^−^) (**Figure 6 i, i’ and j, j’ solid arrows**).

### 15-plex quantification of immune cells in staged MS lesions by IMC

Following the visualization of markers of interest in tissue sections by IMC, we pre-set thresholds for each marker and analysed combinations of markers that identify the phenotype and functional status of immune cells, as shown in **Figure 7 – Supplement Figure 2**. Focusing on the MS tissue, we assembled these data into heat maps to visualize quantitatively the cellular content of each region of interest (**Figure 7a**).

#### Macrophages and microglia

CD45^high^HLA^+^TMEM^−^ macrophages were found in the normal-appearing white matter and a low density of macrophages (3 cells/mm^2^) contained PLP within their CD68^+^ lysosomes, indicative of demyelinating activity. The number of demyelinating CD45^high^HLA^+^TMEM^−^CD68^+^PLP^+^ macrophages drastically increased in the (p)reactive lesions (34.8 cells/mm^2^), reaching peak density in the core of active lesions (36.5 cell/mm^2^) and edge of active-inactive demyelinating lesions (50.5 cell/mm^2^). In the active core we also found a high density (92.6 cell/mm^2^) of CD45^high^HLA^+^TMEM^−^CD68^+^PLP^−^ macrophages, which represent phagocytes with enlarged but empty vacuoles.

Similarly to demyelinating CD45^high^HLA^+^TMEM^−^CD68^+^PLP^+^ macrophages, demyelinating CD45^low^HLA^+^TMEM^+^CD68^+^PLP^+^ microglia were found in high numbers in the (p)reactive lesions with peak density in the core of active lesions (249.5 cell/mm^2^) and edge of active-inactive demyelinating lesions (403.3 cell/mm^2^). Also in line with the distribution of non-demyelinating macrophages, a high density (224.9 cell/mm^2^) of non-demyelianting CD45^low^HLA^+^TMEM^+^CD68^+^PLP^−^ microglia with empty vacuoles were found in the core of active lesions. Overall, we found that in the core of active lesions on average 79% of HLA^+^ cells are microglia and that they constitute 87% of the actively demyelinating (PLP^+^) phagocytes. In the edge of a mixed active-inactive demyelinating lesion, we found that on average 88% of HLA^+^ cells are microglia and that they constitute 89% of actively demyelinating (PLP^+^) phagocytes.

#### T cells

Both CD45^high^CD3^+^CD8α^+^CD4^−^ (CD8) T cells and CD45^high^CD3^+^CD8α^−^CD4^+^ (CD4) T cells were abundant in the MS tissue from the (p)reactive lesional stage (CD8, 53.4 cells/mm^2^, CD4, 49.7 cells/mm^2^) with peak densities in the core of active lesions (CD8, 58.6 cells/mm^2^, CD4, 38.5 cells/mm^2^), the periplaque (CD8, 92.3 cells/mm^2^, CD4, 32.2 cells/mm^2^) and the rim (CD8, 43.2 cells/mm^2^, CD4, 68.8 cells/mm^2^) of mixed active-inactive lesions. Overall, we found that in the core of active lesions on average 60% of T cells are CD8^+^, 3% of which are activated and proliferating (NFAT^+^Ki67^+^). On the contrary, in the edge of a mixed active-inactive lesion, we found that on average 61% of T cells are CD4^+^, 17% of which are activated and proliferating (NFAT^+^Ki67^+^). We verified these findings by examining an independent combination of markers – co-expression of CD38 and HLA on both CD4+ and CD8+ T cells is associated with T cell activation in the context of viral infection[43]. We found that CD4^+^CD38^+^HLA^+^ and CD8^+^CD38^+^HLA^+^ were likewise enriched in the core of the active lesion and the edge of the active/inactive lesion with CD4^+^CD38^+^HLA^+^ T cells being particularly represented at the edge of the active/inactive lesion. However unlike the NFAT^+^Ki67^+^ T cells, CD38^+^HLA^+^ T cells were present in particularly high density in the (p)reactive lesion.

#### B cells

Using the CD38 marker, we were able to further define B cells sub-populations beyond the qualitative images in **Figure 6**. We found B cells across all lesion types with switched memory CD45^high^Igκ/Igƛ^+^IgM^−^CD38^+^ B cells predominating in the core of active lesions (29.2 cells/mm^2^, 58% of all detected B cells) and periplaque white matter (22.2 cells/mm^2^, 87% of all detected B cells) and at the lesion rim (44.0 cells/mm^2^, 44% of all detected B cells) of mixed active inactive lesions.

#### Analysis of the distribution of immune cells in staged MS lesions by IMC

Since the distribution of blood-derived immune cells in relation to blood vessels can inform on the relationship between immune infiltrates and tissue injury, we performed a morphometric analysis of the distance between functional cell types and blood vessels in different MS lesion areas (**Figure 7b**).

#### Macrophages

We found that demyelinating CD45^high^HLA^+^TMEM^−^CD68^+^PLP^+^ macrophages infiltrated the lesion parenchyma in (p)reactive lesions (average distance from blood vessels, 136μm) and periplaque white matter (average distance from blood vessels, 131-191μm), indicating that demyelinating events occur already in tissue that does not show obvious signs of demyelination. Demyelinating macrophages were mostly found in close proximity to blood vessels in active demyelinating lesions (average distance from blood vessels, 87μm) and at the edge of active-inactive demyelinating lesions (average distance from blood vessels, 70μm). Non-demyelinating CD45^high^HLA^+^TMEM^−^CD68^+^PLP^−^ macrophages were found within the lesion parenchyma in both active lesions (average distance from blood vessels, 107μm) and active-inactive lesions (edge: average distance from blood vessels, 244μm; core:average distance from blood vessels, 279μm), representing phagocytes that are no longer actively demyelinating.

#### T cells

In (p)reactive and periplaque white matter, both CD8^+^ and CD4^+^ T cells infiltrated the parenchyma (CD8^+^ T cells: average distance from blood vessels, 114-238μm; CD4^+^ T cells: average distance from blood vessels, 80-208μm). At the edge (CD8^+^ T cells: average distance from blood vessels, 45μm; CD4^+^ T cells: average distance from blood vessels, 21μm) and core (CD8^+^ T cells: average distance from blood vessels, 121μm; CD4^+^ T cells: average distance from blood vessels, 89μm) of active-inactive lesions, CD4^+^ T cells were located in closer proximity to blood vessels compared to CD8^+^ cells, which instead appeared to diffusely infiltrate the lesional parenchyma. CD8^+^ and CD4^+^ T cells were found to equally infiltrate the parenchyma in active lesions (CD8^+^ T cells: average distance from blood vessels, 89μm; CD4^+^ T cells: average distance from blood vessels, 88μm).

#### B cells

We found that naïve CD45^high^Igκ/Igƛ^+^IgM^+^CD38^−^ B cells and switched memory CD45^high^Igκ/Igƛ^+^IgM^−^CD38^+^ B cells infiltrated the parenchyma in both (p)reactive lesions (naïve B cells: average distance from blood vessels, 62μm; memory switched B cells: average distance from blood vessels, 92μm) and periplaque white matter (naïve B cells: average distance from blood vessels, 71μm; memory switched B cells: average distance from blood vessels, 103-113μm). Within lesions, naïve B cells were focally located in the perivascular space of veins at the the edge (average distance from blood vessels, 2μm) and core (average distance from blood vessels, 9μm) of active-inactive lesions. Switched memory CD45^high^Igκ/Igƛ^+^IgM^−^CD38^+^ B cells were present in the vicinity of blood vessels at the rim (average distance from blood vessels, 44μm) and core (average distance from blood vessels, 19μm) of active-inactive lesions but were found to also diffusely infiltrate the parenchyma of active lesions (average distance from blood vessels, 82μm).

## DISCUSSION

In this study we used imaging mass cytometry to multiplex 15+ markers to stain a single tissue section. The panel contained both cell-specific and functional markers, and enabled the analysis of single-cell phenotypes and functional states of resident microglia, blood-derived (recruited) macrophages, T and B lymphocytes in demyelinating and highly inflammatory lesions in a case of severe rebound MS disease activity after natalizumab cessation[26]. We first showed the validity of the technology on post-mortem MS brain tissue and then applied it to the analysis of immune cells in the lesions compared to control brain tissue.

IMC reproduced IHC- and IF-equivalent staining patterns with no apparent changes in specificity compared to standard IF. Therefore, antibodies validated with IF for the study of the MS brain will likely be applicable to the IMC approach. It should be noted, however, that the concentration and the staining conditions of some IHC- osr IF-verified antibodies may not be implemented as is into the IMC protocol: titration and/or amplification (for example with biotin-streptavin) of pathologist-verified antibodies is required for optimal visualization by IMC.

In addition to visualizing a multitude of cell types, IMC allows for inclusion and exclusion criteria of selected markers to provide better confidence of cell identity. Furthermore, the highly quantitative nature of the IMC approach enables the analyses of data with pre-set thresholds for each marker, and permits further validation based on combination (inclusion/exclusion) with other markers. For example, we were able to distinguish CD45^high^ cells that were TMEM119^−^CD68^+^ thus identifying macrophages *versus* CD45^low/+^ cells that were TMEM119^+^CD68^+^ thus identifying microglia.

We found that in the control brain, microglial cells lose their homeostatic phenotype and acquire an activated state. This is in line with an earlier study demonstrating expression of certain activation markers by microglia within the normal human brain, and it is in agreement with recent immunohistological findings that show no expression of the homeostatic molecule P2RY12[42] in 48% of microglia in control brains[44]. Whether this activation state is the result of systemic exposure to recurrent infections[33] or is the result of vascular and neurodegenerative changes related to normal ageing[11] (the control subject was 86 years), or whether it is an inherent property of microglia in the human brain[44], is unclear.

In line with recent observations in carefully staged lesions from a large cohort of MS patients at well-defined disease stages[44], we found that microglial activation was not restricted to lesional tissue but was also present in the normal-appearing white matter and (p)reactive lesion site. In these regions, although myelination appeared normal, we also found that demyelinating blood-derived macrophages infiltrated the parenchyma. In active lesions and in the active edge of mixed active-inactive demyelinating lesions[25] (slowly expanding or ‘smouldering’[15]), microglia and macrophages displayed similar phenotypic changes characterized by the predominant expression of markers associated with activation and phagocytosis. In contrast, in the core of mixed active-inactive demyelinating lesions, microglia and macrophages lost expression of molecules involved in antigen presentation and drastically reduced their phagocytic activity, as previously described[44]. Notably, a large proportion (on average 88%) of demyelinating macrophage-like cells in active lesions and at the edge of mixed active-inactive lesions were derived from the resident microglial pool, whereas macrophages that infiltrated the parenchyma of these lesional areas were largely inactive as indicated by the presence of enlarged but empty vacuoles in these cells. This is likely the result of the macrophage’s inability to digest the myelin’s neutral lipid components that accumulate and persist in macrophages.

In terms of lymphocytes, in classical active lesions and mixed active-inactive demyelinating lesions we showed that T cells were abundant. Although CD8^+^ T cells generally predominated across lesional stages, and in some cases proliferated (as also shown in a recent study[31]), interestingly we found a conspicuous number of CD4^+^ T cells not only within lesions but also at the (p)reactive lesion site and periplaque white matter. In addition, CD4^+^CD38^+^HLA^+^ “chronically activated” T cells[43] were also particularly abundant in the (p)reactive lesion site. This suggests an involvement of these cells in the early stages of lesion formation, even in established lesions. Similar to other findings in the case of T lymphocytes, our data also reproduced immunohistological findings that described B lymphocytes in all lesion stages in lower numbers compared to T cells [31]. By using IgM in combination with CD38 and *κ/λ* our panel has the increased capacity of identifying different B cell subsets. Indeed, by using IgM in combination with CD38 and *κ/λ*, our panel has revealed different B cell subsets including IgM^+^ and switched memory B cells.

With the IMC approach, we were able to reproduce findings by Machado-Santos et al wherein they proposed that although active demyelination is associated with activated blood-derived macrophages, it is largely driven by the resident activated microglial pool[44]. This was also suggested by earlier studies[27], pointing to the possibility that therapeutic intervetions aimed at blocking entry of myeloid cells from the circulation into the brain parenchyma may be insufficient to halt the disease process.

It has been suggested that CD8^+^ T cells in lesions from patients with relapsing, progressive and fulminant acute MS show features of tissue-resident memory cells and play a central role in the establishment of tissue-specific immunological memory, propagating chronic compartmentalized inflammation and tissue damage in the MS brain by local activation following re-exposure to their cognate antigen[31]. B cells are also detected in all MS lesion types but their localization seems to be restricted to the pervascular space of some veins[31]. Machado-Santos et al have shown that the majority of B and T cells are present in the perivascular cuffs, distant from sites of initial myelin damage[31]. This supports the possibility that demyelination is induced by soluble factors produced by lymphocyte which diffuse into the tissue and in turn activate phagocytes. Our findings from a single MS case with high inflammatory activity only incompletely reproduce these findings. While we also observed perivascular localization of B and T cells, these lymphocytes were also found to diffusely infiltrate the lesion parenchyma, which could support a contact-mediated active demyelination, at least in this MS case. However, due to the nature of the acquired region of interest, which doesn’t capture the areas surrounding the site of ablation, it is possible that blood vessels were positioned immediately outside the region of interest. These would be missed in the cell-blood vessel distance analysis, which would result in considering cells that may in reality have a perivascular localization, as been located far from blood vessels. Further analysis of larger areas across multiple tissue samples are required to better answer this question.

A comprehensive phenotypic characterization of B cells in MS tissue is lacking and the role of B cells in MS lesions is currently unresolved. Recent clinical studies have reported a protective effect of therapies targeting CD20^+^ B cells in MS patients, suggesting a major role for B cells in the disease process[20,21]. Our IMC results allowed for better segregation of B cell phenotypes (memory, class switched etc). Further addition of other markers, particularly for plasma cells (such as CD138, TACI) will be important for a full characterization of B cell subsets within the MS brain. This is particularly relevant in light of recent findings that demonstrate that some B lineage cells play a protective role in neuroinflammatory processes[37].

While IMC has the advantage of multiplexing capability, it also has limitations. For example, it yields information only about the brain region imaged and is low throughput. It is therefore possible that different cell populations can exist in brain regions and sublesional areas other than those imaged. As is the case for the analysis of tissue stained with standard IHC/IF methods, multiple regions must be acquired. Another limitation, that seems apparent from the tissue that we examined, is that cell densities by IMC were higher than those derived by IF (**Figure 1**). This may be due to the fact that we had to use a serial section to compare the two methods, with the two sections containing slightly different numbers of cells. Alternatively, IMC may have an increased sensitivity in brain tissue compared to IF. However, in spite of these differences, the proportionality of the output signal for IMC vs IF was consistent across samples (Figure 1 and [9]). In addition, our study has the limitation that it is based on the analysis of immune cells in lesions from a single MS. Our goal was not to uncover novel cell populations in the MS brain, but rather to provide proof that the IMC technology can be used as a powerful tool for the analysis of complex cellular phenotypes in heterogeneous tissues such as the MS brain. Moreover, given that the cells in these lesions had known phenotypes, the supervised approach for thresholding used herein was reasonable. However in the future, unsupervised analysis of data sets generated using the IMC approach may identify novel cell types in tissue that is understudied, for example the MS meninges. In addition, discovery of novel cell types using a technique such as IMC can then be recapitulated with standard techniques using multiple well-characterized specimens from established brain banks.

Overall, our data reproduced immunohistological patterns of microglia and lymphocytes activation described in carefully staged MS brain lesions at well-defined disease stages[31,44] using a multi-parameter approach. The significance of observed B cells of IgM memory and class switch memory phenotypes, warrant further study. We propose that IMC will enable a high dimentional analysis of single-cell phenotypes along with their functional states, as well as cell-cell interactions in relation to lesion morphometry and (demyelinating) activity. The IMC approach in combination with exhisting imaging techniques, can profoundly impact our knowledge of the nature of the inflammatory response and tissue injury in the multiple sclerosis brain.

## Acknowledgements

We would like to thank the brain donors.

This work was funded by a team grant from the MS Society Research Foundation to AP and JG, and the National Multiple Sclerosis Society Research Grant RR-1602-07777 to VR.

## Author contributions

V.R. performed the pathological characterization for the MS lesions, guided the acquisition of the IMC data and contributed to data analysis. S.S.M. performed the IF and IMC staining, and contributed to the data analysis. K.L. contributed to the optimization of the IMC staining. O.L.R and V.R. supervised the optimization of the IMC staining. S.Z. performed the histological staining. A.P. acquired the patient tissue. Both A.P. and O.O. contributed intellectually. E.S. acquired the IMC data. T.D.M. and F.F. analysed the data. J.L.G. supervised the study. V.R and J.L.G. contributed to the design of the study and wrote the manuscript.

## Figure Supplements

**Figure 1 – Figure Supplement 1.**
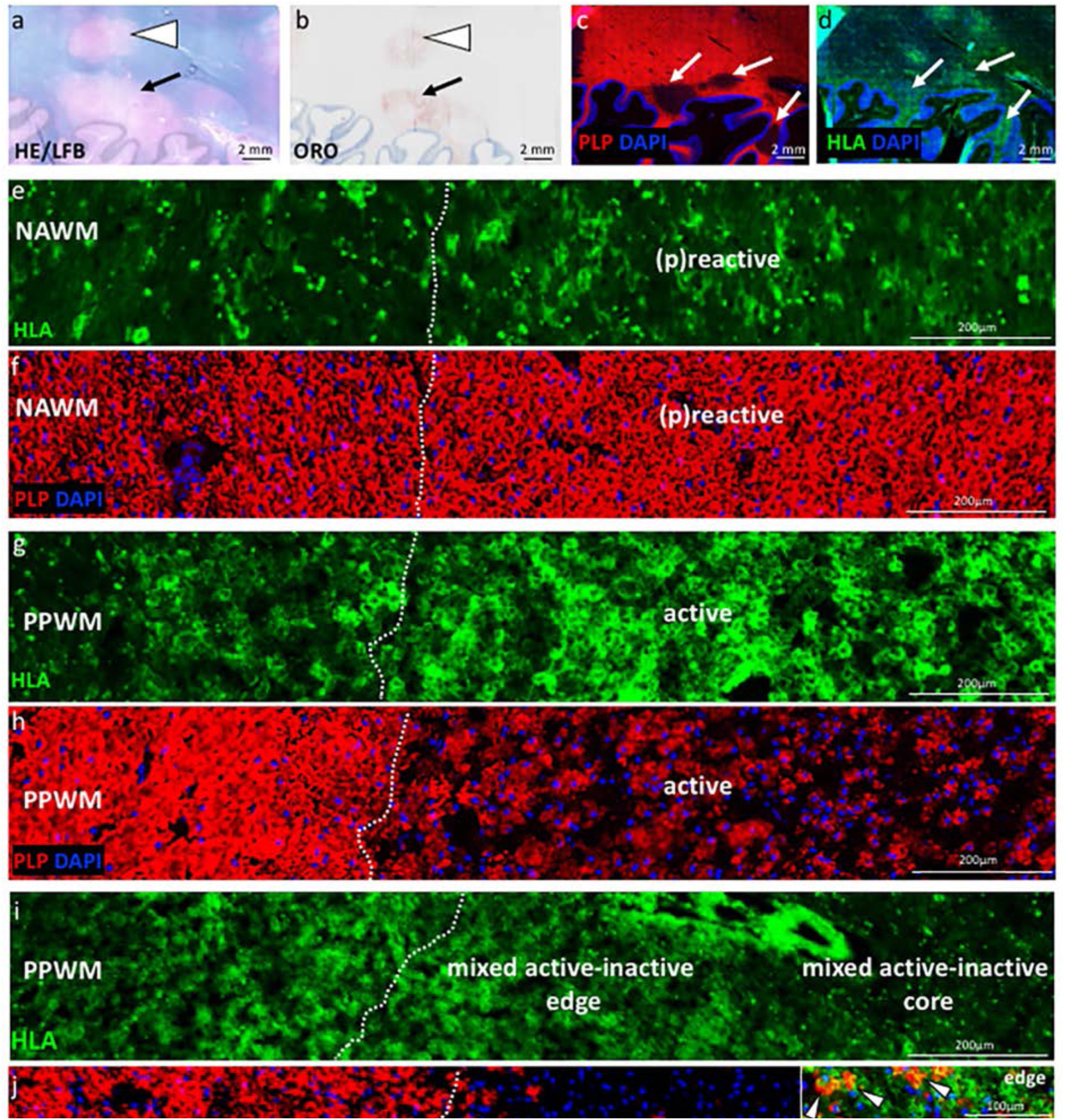
Staging of MS lesions by IF. General pathology: demyelinating lesions (arrows and arrows head in **a** and **b**) seen in hematoxylin & eosin (HE)/Luxol fast blue (LFB) stain of myelin (**a**) and oil red o (ORO) stain of neutral lipids within macrophages (**b**). Lesional pathology: demyelinating lesions (arrow heads in **c** and **d**) visualized by proteolipid protein (PLP in red) stain of myelin (**c**) and human leukocyte antigen (HLA, in green) stain of microglia/macrophages (**d**). (**e-j**) Low magnification images of HLA and PLP stains, depicting the distribution and morphology of HLA+ microglia/macrophages and myelin in different sites and lesion stages. (**e-f**) (P)reactive lesion (block no. CR4A): Note the increase in microglia/macrophage reactivity at the (p)reactive lesion site compared to the normal-appearing white matter (NAWM) (**e**), with normal PLP myelin stain seen across the NAWM and (p)reactive lesion (**f**). (**g-h**) Active demyelinating lesion (block no. CR4A): low glia reactivity and normal myelin stain is seen in the periplaque white matter (PPWM). Profound microglia/macrophage activation is seen in the active lesion (**g**), where myelin is being destroyed (**h**). (**i-j**) Mixed active/inactive demyelinating lesion (block no. CL3A): low glia reactivity and normal myelin stain is seen in the periplaque white matter (PPWM). An increased density of HLA^+^ cells with the morphology of microglia/macrophages is seen at the active SEL edge, with degraded PLP^+^ myelin within macrophages (arrows head in inset). In contrast, there are only few HLA^+^ cells at the inactive lesion center.

**Figure 1 – Figure Supplement 2.**
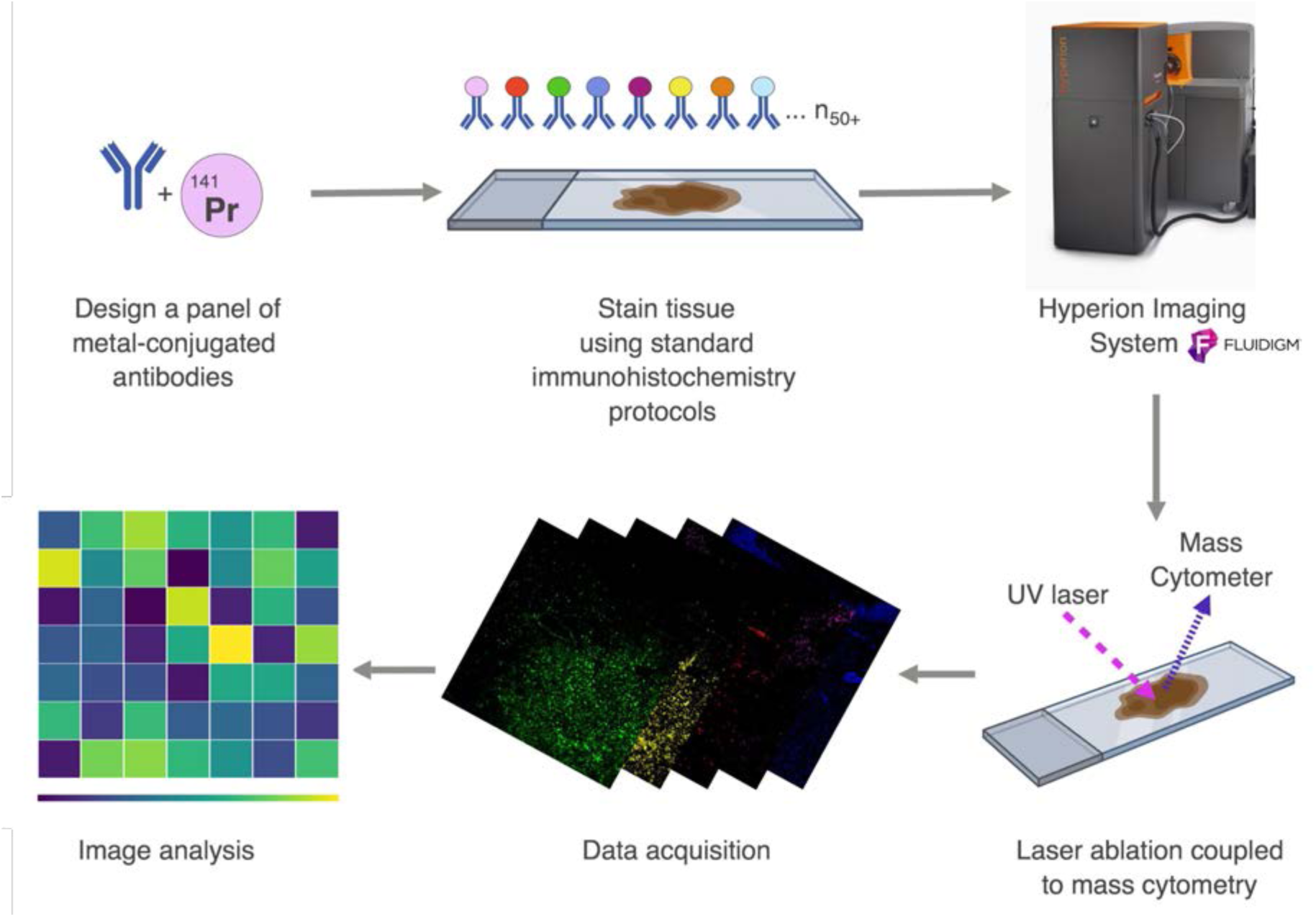
Workflow of Imaging Mass Cytometry. A panel is designed using pathologist-verified antibodies conjugated to metals. The brain tissue is stained simultaneously with a cocktail of all the metal-conjugated antibodies and placed into the Hyperion Imaging System. The tissue is ablated by a UV laser beam (λ = 219 nm). A plume of particles produced by the laser is taken up by a flow of inert helium or argon gas and introduced into the CyTOF mass cytometer (Hyperion Imaging System from Fluidigm (formerly DVS Sciences)). Isotopes associated with each spot are detected and indexed against the source location, yielding an intensity map of the target proteins throughout the tissue, creating spatially resolved images of multiple parameters. The acquired data is analyzed and visualized using heat maps.

**Figure 1 – Figure Supplement 3.**
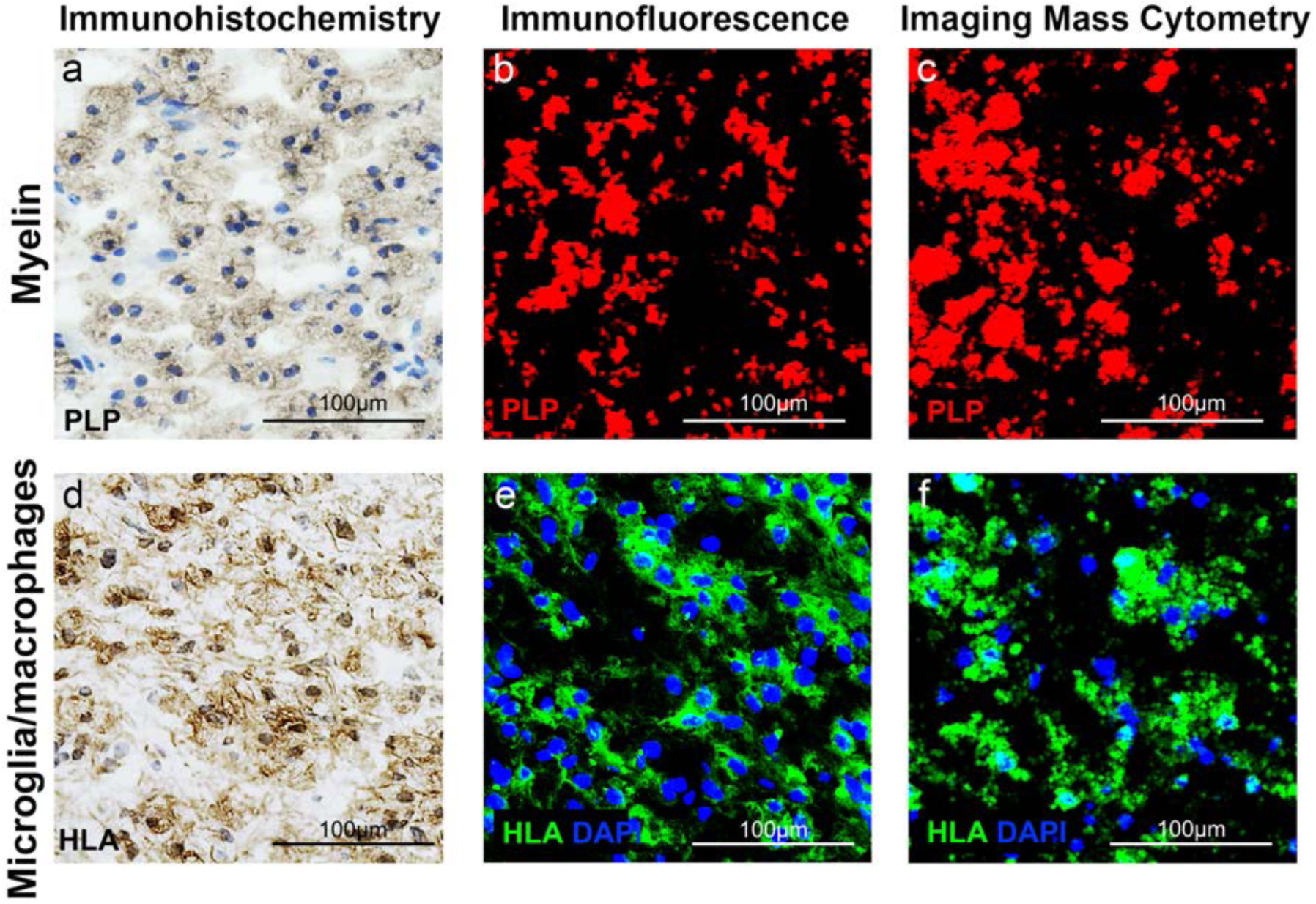
Validation of IMC staining patterns in MS lesions. Core of an active demyelinating lesion showing reduced proteolipid protein (PLP) stain by immunohistochemistry (**a**), immunofluorescence (**b**) and IMC (**c**) and corresponding areas stained with anti-HLA to detect antigen presenting cells by immunohistochemistry (**d**), immunofluorescence (**e**) and IMC (**f**).

**Figure 7 – Figure Supplement 1.**
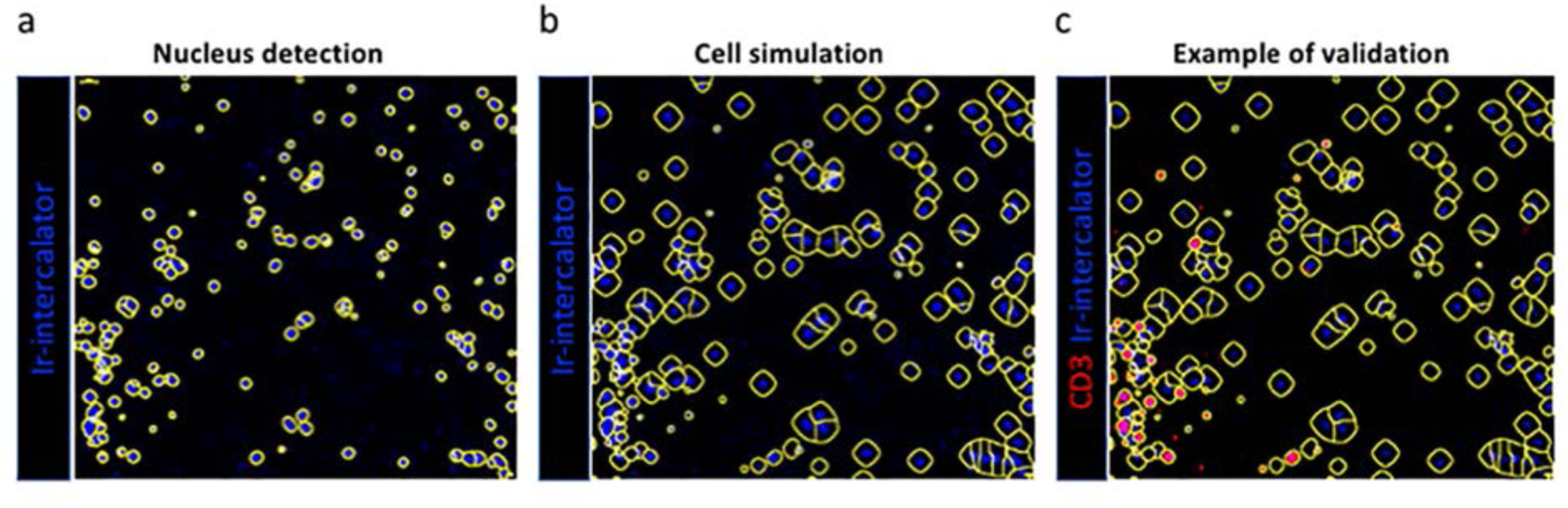
Single cell segmentation. A Gaussian blur was applied to the DNA signal (nucleus detection - **a**), and the resulting blurred image was segmented to identify nuclear content corresponding to individual cell areas using a combination of threshold and watershed filters (cell simulation - **b**). Subsequently, we interrogated the segmented image for the presence of specific markers or combinations of markers that are either biologically co-expressed, or whose expression is mutually exclusive. In this example we show CD3 (example of validation - **c**).

**Figure 7 – Figure Supplement 2.**
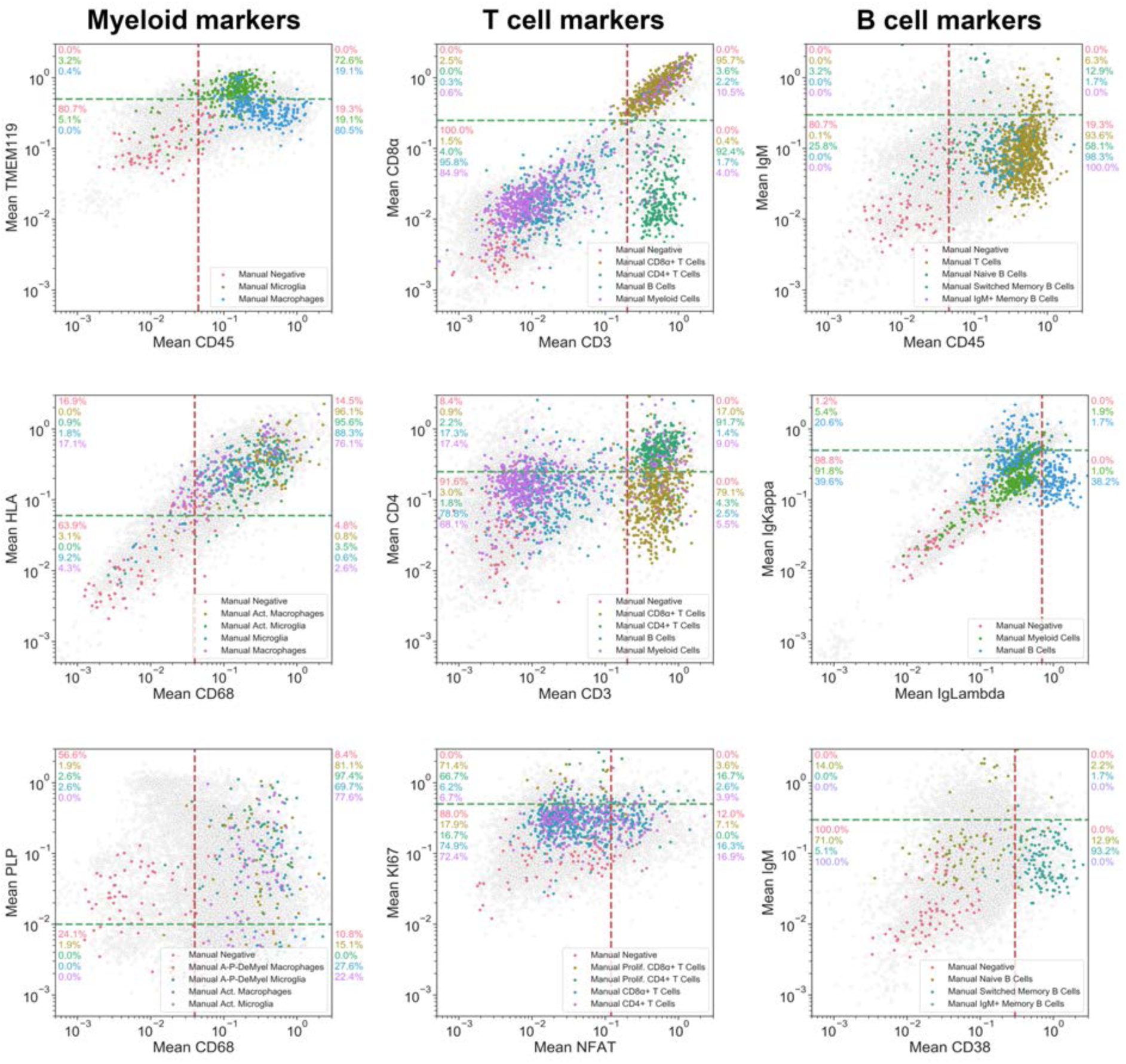
Gating strategy used for the identification of cell subsets. Gating strategy for the identification of cell subset phenotypes and activation states of microglia, macrophages, T cells and B cells. In brief, the per-cell mean intensities of specific marker combinations are shown here in 2D log-log biaxial scatterplots. Gates were established based on pathologist-verified positive cells (see coloured cells superimposed into each dotplot contrasting with non-verified cells in grey).

**Figure 7 – Figure Supplement 3.**
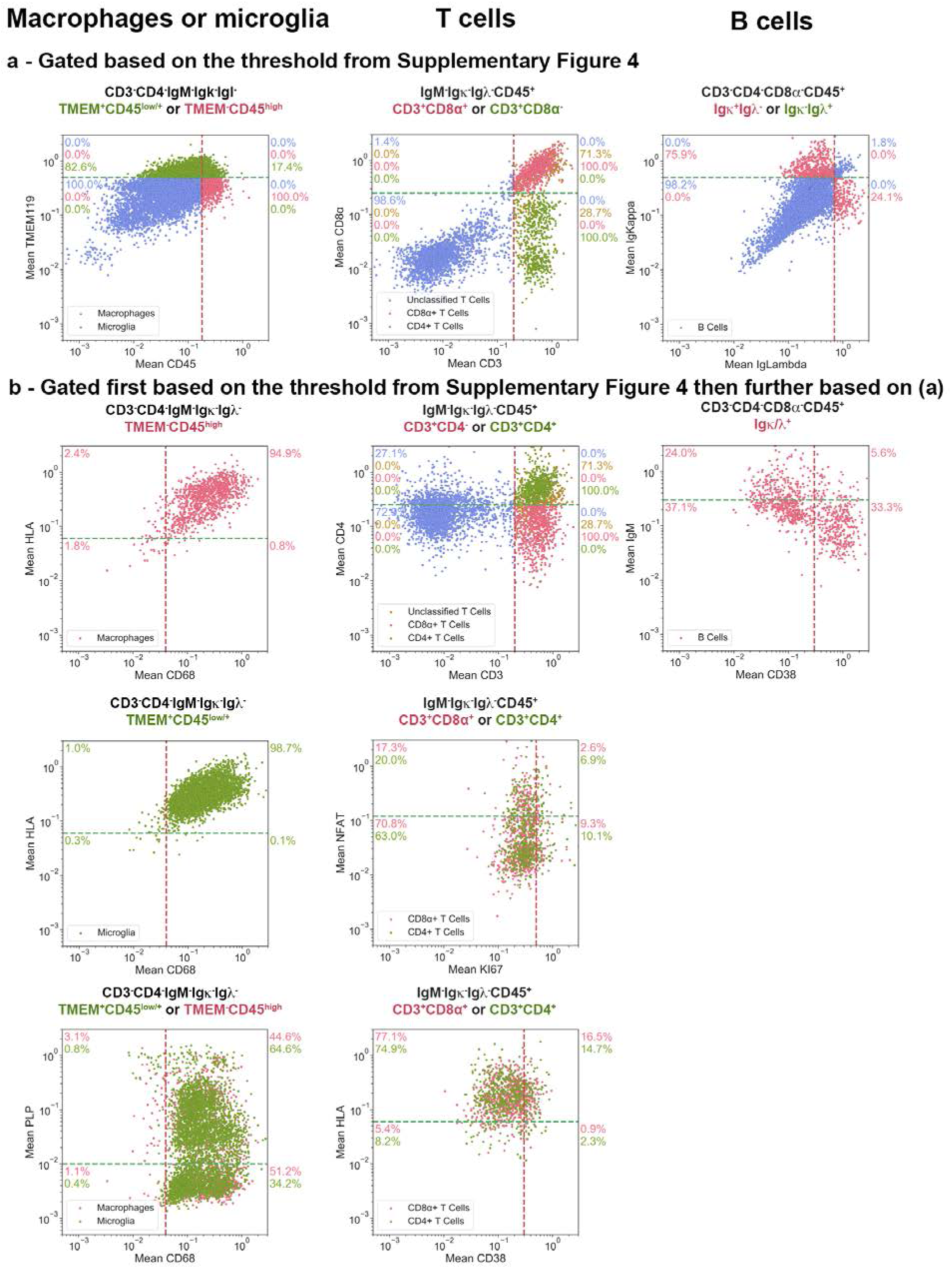
Gating strategy used for the generation of heat maps. Using the quadrants that capture the appropriate positivity range of each cell phenotype shown in **Figure 7 – Figure Supplement 2**, cells were subjected to the positive and negative gating strategies as outlined in the Materials and Methods for each lineage and indicated in (**a**). Subsequently, these cells were plotted for the marker combinations listed in **Table 3**. The frequency of cells in each quadrant are indicated. Note that some CD3^+^CD45^+^ T cells could not be classified because they fell outside of the specified gates for either of the two markers – CD8^+^ cells that were not simultaneously CD4^−^, or CD4^+^ cells that were not simultaneously CD8^−^. This is due to the dynamic range of these particular markers and thus our inability to get a clean CD4^+^CD8^−^ or CD4^−^CD8^+^ T cell population. Cells that fulfilled the gating criteria specified above each image, but which did not fulfill the requirements for classification as Macrophages, Microglia, B cells or T cells, are shown in blue.

## Source data

**Figure 7 – Source data 1**

Source file for quantitative data of all ROI:

ROI 2 from block 95-056 (white matter control)

ROI 1 from block CL3a (active lesion)

ROI 2.1 from block CL3a (mixed active-inactive lesion)

ROI 2.2 from block CL3a (mixed active-inactive lesion)

ROI 3 from block CL3a (mixed active-inactive lesion)

ROI 4 from block CL3a (active lesion)

ROI 2 from block CR4a (mixed active-inactive lesion)

ROI 4 from block CR4a (active lesion)

ROI 8 from block CR4a (normal-appearing white matter)

ROI 3 from block CR4a (active lesion)

ROI 1 from block CR4a ((p)reactive lesion)

ROI 5 from block CR4a (active lesion)

ROI 6 from block CR4a (mixed active-inactive lesion)

